# Coiled-coil and RPW8-type immune receptors function at the plasma membrane in a phospholipid dependent manner

**DOI:** 10.1101/2020.11.18.388520

**Authors:** Svenja C. Saile, Frank M. Ackermann, Sruthi Sunil, Adam Bayless, Eva Stöbbe, Vera Bonardi, Li Wan, Mehdi Doumane, Yvon Jaillais, Marie-Cécile Caillaud, Jeffery L. Dangl, Marc T. Nishimura, Farid El Kasmi

**Author notes:** Novozymes North America Inc, 108 T W Alexander Drive Bldg 1 A, Raleigh, NC, United States of America. CAS Center for Excellence in Molecular Plant Sciences / Institute of Plant Physiology and Ecology, 300 Feng Lin Road, Shanghai 200032, China. These authors contributed equally: Saile, S.C. and Ackermann, F.M.

## Abstract

Activation of intracellular nucleotide-binding leucine-rich repeat receptors (NLRs) results in immunity and a localized cell death response of infected cells. Cell death activity of many NLRs requires oligomerization and in some cases plasma membrane (PM) localization. However, the exact mechanisms underlying PM localization of NLRs lacking recognizable N- or C-terminal lipidation motifs or predicted transmembrane domains remains elusive. Here we show that the PM localization and stability of members of the RPW8-like coiled-coil (CC_R_) domain NLRs (RNLs) and a CC-type NLR (CNL) depend on the interaction with PM phospholipids. Depletion of phosphatidylinositol-4-phosphate (PI4P) from the PM led to a mislocalization of the analyzed NLRs and consequently inhibited their cell death activity. We further demonstrate activation-dependent self-association of cell death inducing RNLs. Our results provide new insights into the molecular mechanism of NLR PM localization and defines an important role of phospholipids for CNL and RNL activity during immunity.

## Introduction

Plant intracellular immune receptors of the nucleotide-binding leucine-rich repeat receptor (NLR) family mediate recognition of pathogen-derived effector proteins and the induction of a strong immune response. In many cases, NLR activation leads to the hypersensitive response, a type of programmed cell death of the infected cells^1–3^. Based on their N-terminal domain architecture, three classes of NLRs have been described in plants: Toll/Interleukin-1 receptor (TIR) NLRs (TNLs), coiled-coil (CC) NLRs (CNLs) and the RPW8-like coiled-coil (CC_R_) domain NLRs (RNLs)^1^. In *Arabidopsis thaliana* (Arabidopsis) the RNL subclass consists of two gene families, *ACTIVATED DISEASE RESISTANCE 1* (*ADR1*) and *N REQUIREMENT GENE 1* (*NRG1*), both being required for immune signalling and cell death induction of many other NLRs, particularly TNLs^4–9^. CNLs, TNLs and most likely RNLs induce immune signalling and cell death by oligomerization^10–13^. CNL activation is speculated to result in the formation of a pore-like or membrane disrupting structure of the CC domain (a so-called resistosome) at the plasma membrane (PM)^10,14–16^. PM localization was demonstrated to be required for cell death and immune function of many CNLs, including Arabidopsis RPS5, RPM1 and ZAR1^10,17–20^. The subcellular localization of RNLs has not yet been analysed in detail. So far only a potential endoplasmic reticulum (ER) localization as well as a partial PM localization of AtNRG1s was described^6,7^. Interestingly, many PM-localized CNLs and the RNLs have no predicted N- or C-terminal lipidation motif or transmembrane domain/sequence and the mechanism that tethers them to the PM is unknown^17^. Thus, the molecular determinants driving their localization and cell death function at the membrane are not identified.

Homology modelling suggests that the CCR domains of RNLs share structural similarities with the N-terminal 4-helix bundle (HeLo domain) of mammalian mixed-linage kinase domain-like (MLKL) proteins and fungal HET-s/HELL proteins^21–23^. HeLo domains mediate the cell death function of MLKL and HET-s/HELL proteins and are proposed to oligomerize and disrupt or permeabilize the PM^24,25^. Cell death function and PM localization of MLKL proteins requires the interaction of their HeLo domain with specific phospholipids at the PM^26,27^.

Negatively charged, anionic phospholipids at membranes mediate electrostatic interactions with many proteins that contain polybasic, basic hydrophobic or cationic domains or clusters^28,29^. Phosphatidylinositol-4-phosphate (PI4P) is one of the major phospholipids of the plant PM and a main driver of PM electronegativity^30^. Expression of the PM-anchored catalytic domain of the yeast phospholipid-phosphatase Sac1p protein, which specifically dephosphorylates PI4P and therefore reduces PI4P levels and the PM electronegativity^30,31^, can be used to determine whether a protein requires the presence of PI4P (or a high electronegativity) for localization and/or function at the PM. Depletion of PI4P from the PM affects the localization and function of several proteins, including the auxin transport regulator PINOID or the BRI1 kinase inhibitor 1, BKI1^30^.

We show that decreasing PI4P abundance at the PM results in mis-localization and rapid degradation of AtRPM1 (a CNL) and the three AtADR1s (RNLs). Further, depletion of PI4P also severely affected cell death induction mediated by AtRPM1, as well as both AtRNL subfamilies.

Our results provide new insights into the molecular mechanism of NLR PM localization and defines an important role of the PM phosphatidylinositol phosphate (PIP) pool in CNL- and RNL-mediated cell death induction during plant immunity. Further, our work indicates that CNL- and RNLs deploy a lipid-protein interaction similar to animal MLKL proteins for PM localization, which is likely necessary for cell death execution at the PM.

## Results

### Arabidopsis ADR1s localize at the plasma membrane in *Nicotiana benthamiana*

The tentative subcellular localization of two Arabidopsis full length RNLs, AtNRG1.1 and AtNRG1.2, was recently described. Both proteins localize at ER membranes, partially at the PM and potentially in the cytosol when transiently over-expressed in *Nicotiana benthamiana* (*N. benthamiana*) and measured by confocal microscopy^6,7^. Their intracellular localization was not changed upon effector-triggered and TNL-mediated activation^6,8^. However, there is no information on the localization of either pre- or post-activated AtADR1s, the other RNL subfamily. To investigate the subcellular localization of the three AtADR1 proteins pre- and post-activation we transiently expressed C-terminally EYFP- or Citrine-HA-tagged wildtype ADR1, ADR1-L1 and ADR1-L2 in *N. benthamiana* leaves. We observed a strong co-localization of all three wildtype AtADR1s with the PM-localized receptor-like kinase BRI1-mRFP (Fig. 1a,c,e; Supplementary Fig. S1a,e,g)^32^. In contrast to AtADR1-L1 and AtADR1-L2 the localization of AtADR1 was not restricted to the PM. AtADR1 additionally localized to some puncta, of which some might be PM and/or ER associated (Supplementary Fig. S1a and c), and the ER membrane, where it co-localized with the ER-resident plant V-ATPase assembly factor AtVMA12-RFP (Supplementary Fig. S1c)^33^.

**Fig.1.**
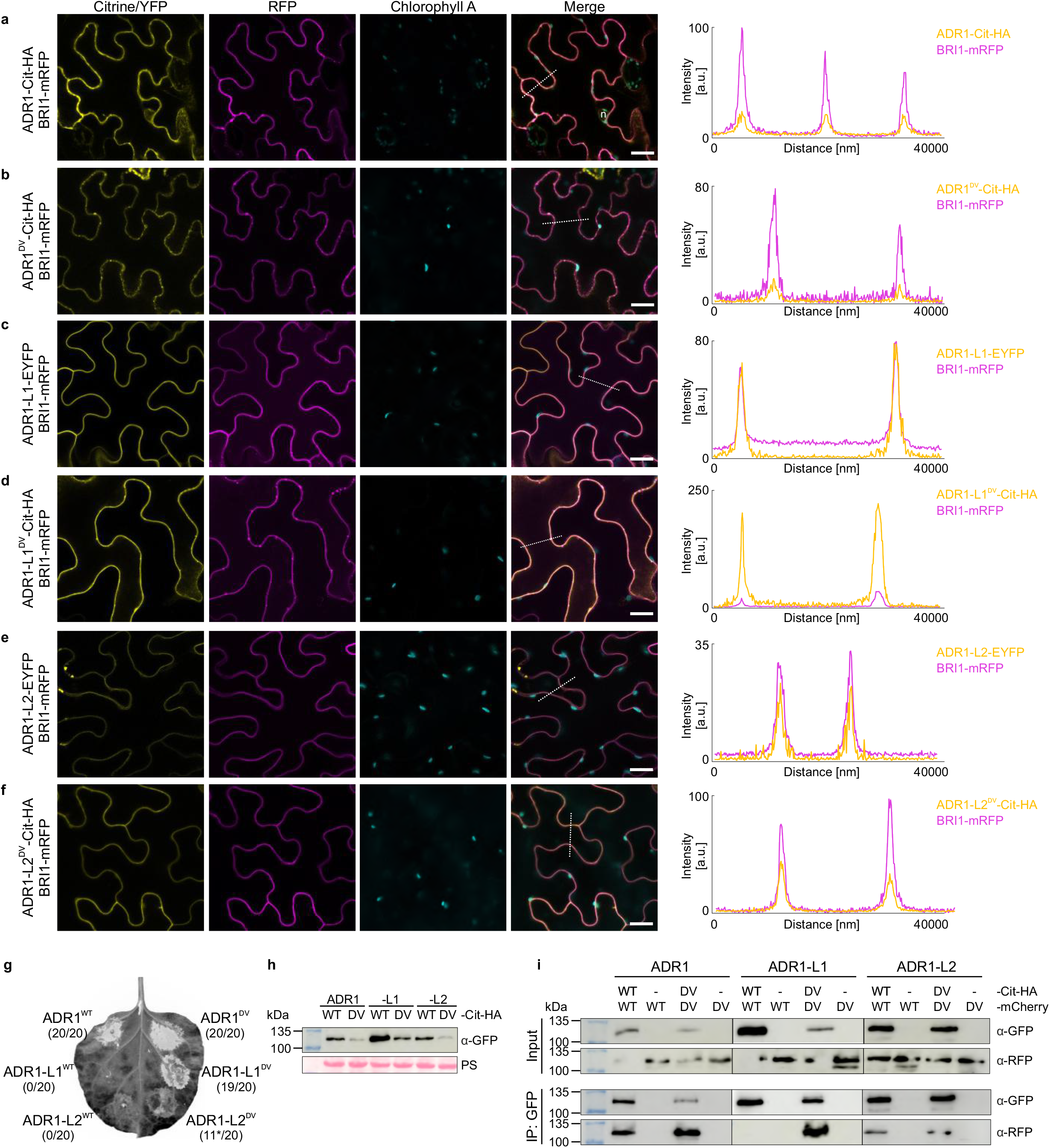
AtADR1 proteins mainly localize to the PM and do self-associate. **a-f,** Single plane secant views showing AtADR1 proteins (ADR1, ADR1-L1, ADR1-L2) localize at the plasma membrane (PM). The indicated ADR1 proteins fused to Citrine-HA or EYFP were transiently co-expressed with the PM-resident protein BRI1-mRFP in *N. benthamiana* leaves and confocal imaging was done at 4 (a, b, d) or 5 hours (f) post Estradiol induction or 2 days post infiltration (c, e). Localization of ADR1s is shown with the first column (Citrine/YFP, in yellow) and the co-localized PM-localized BRI1 is shown in the second column (RFP, in magenta). Chloroplasts are shown in the third column (Chlropohyll A, in cyan) and the merged images are shown in the fourth column (merge). Fluorescence intensities were measured along the dotted line depicted in the merge images. Scale bars, 20 μm. **g-i,** ADR1s cell death function coincides with strong protein self-association. **g,** Transient expression of steady-state (WT) or mutant auto-activated (DV) ADR1 s-Citrine-HA fusion proteins in *N. benthamiana*. Photos were taken under UV light at 23 hours post Estradiol induction and 47 hours post infiltration. White areas correspond to dead leaf tissue. Numbers represent the number of leaves showing cell death out of the number of leaves analysed. Asterisk indicates weak cell death. **h,** Immunoblot analysis of the proteins infiltrated in (g) using anti-GFP antibody. Equal loading of the proteins is indicated by the Rubisco band from the Ponceau staining (PS). Proteins were extracted 4 hours post Estradiol induction. **i,** Auto-activated DV mutant ADR1 proteins self-associate. The indicated proteins were transiently co-expressed in *N. benthamiana* and samples were harvested 4 hours post Estradiol induction. Total proteins were immunoprecipitated with anti-GFP beads and immunoblotted with anti-GFP and anti-RFP antibody. Immunoblots using total proteins prior to immunoprecipitation are shown as input (upper panel) and immunoprecipitated proteins are shown in lower panel. Co-immunoprecipitation was repeated three times with similar results. ADR1^DV^: ADR1^D461V^, ADR1-L1^DV^: ADR1-L1^D489V^, ADR1-L2^DV^: ADR1-L2^D484V^.

NLR localization may change when the receptor is activated^10^. To test this, we generated autoactive alleles of ADR1s by mutating a conserved aspartic acid in the MHD motif to valine (QHD to QHV in AtRNLs; cell death phenotype of autoactivated AtRNLs shown in Fig. 1g)^34–36^. The auto-activated AtADR1^DV^ localization appeared more punctate compared to the wildtype AtADR1 (Fig. 1b and Supplementary Fig. S1b and d), indicating that AtADR1 activation could result in a more clustered localization. Similar to AtADR1-L1 and AtADR1-L2 wildtype proteins autoactivated AtADR1-L1^DV^ and AtADR1-L2^DV^ localized to the PM (Fig. 1d,f; Supplementary Fig. S1f,h). We also noticed that AtADR1^DV^ and AtADR1-L1^DV^ localized to BRI1-mRFP positive puncta (Fig. 1b,d; Supplementary Fig. S1b,f), most likely endosomes. This potential endosomal localization was not observed for AtADR1-L2^DV^ (Fig. 1f; Supplementary Fig. S1h).

These results demonstrate that three members of the Arabidopsis ADR1 subfamily localize at the plant PM pre- and post-activation and further suggest that wildtype (steady-state) AtADR1 additionally localizes to ER membranes and ER-associated dot-like structures, as observed for AtNRG1.1^6,7^. The PM localization of the (auto-)activated AtADR1s suggests that they could also execute their immune (cell death) function at the PM.

### Self-association of cell death inducing Arabidopsis ADR1s

NLR function in plants and animals is proposed to require oligomerization for proper induction of cell death and immunity^37^. However, self-association of RNLs was only shown for *N. benthamiana* NRG1^8^. To test whether Arabidopsis ADR1s self-associate and whether self-association is dependent on the activation status, we co-expressed differently tagged wildtype and the QHV (autoactivated) mutant AtADR1s and analysed their self-association by co-immunoprecipitation. We observed that transient over-expression of AtADR1, AtADR1^DV^ and AtADR1-L1^DV^ induced a strong hypersensitive response-like cell death (HR), while the over-expression of ADR1-L2^DV^, although expressed, only resulted in a weak HR that was not reliably reproducible (only 11 of 20 leaves showed HR symptoms; Fig. 1 g and h). We also found that the AtADR1-induced HR occurred earlier in comparison to the AtADR1-L1^DV^ and AtADR1-L2^DV^ triggered HR. Wildtype AtADR1-L1 and At-ADR1-L2 did not trigger a cell death response under our conditions (Fig. 1 g-h). These data are consistent with the hypothesis that wildtype AtADR1 is already highly active under steady-state conditions, whereas AtADR1-L1 and AtADR1-L2 are inactive under steady-state conditions. However, introduction of the D to V mutation in the QHD motif renders AtADR1-L1^DV^ and AtADR1-L2^DV^ into active proteins.

Our co-immunoprecipitation experiments revealed that the proteins inducing a strong cell death response (AtADR1, AtADR1^DV^ and AtADR1-L1^DV^) also strongly self-associated (Fig. 1i). We also observed self-association of wildtype AtADR1-L2 and the autoactivated AtADR1-L2^DV^, however this interaction was much weaker than for the highly active AtADR1, AtADR1^DV^ and AtADR1-L1^DV^ (Fig. 1 i). These results indicate a correlation between self-association and cell death induction, suggesting that the AtADR1s might activate HR via a similar mechanism as canonical CNLs – by the formation of an oligomeric complex at the PM.

### Arabidopsis RNL and CNL PM localization and protein stability requires PM PI4P

The PM localization of AtADR1, AtADR1-L1 and AtADR1-L2 also suggests that they execute their immune function at this cellular compartment as observed for other NLRs, such as AtRPM1^17^. Interestingly, for both RNL families and many PM-localized CNLs, including AtRPM1, no transmembrane region or N- or C-terminal protein lipidation motif could be identified and thus, they are most likely peripheral membrane proteins (Supplementary Table 1)^38^. Given the predicted structural homology of RNL CC_R_ domains with the phosphatidyl-inositol phosphate binding HeLo domain of mammalian MLKL^21^, we investigated whether the presence of specific phosphoinositide species might be important for the PM localization of Arabidopsis RNLs and CNLs. Since PI4P is one of the major phospholipids of the plant PM^30^, we tested whether RNL and CNL PM localization depends on PI4P. Transient expression of catalytic domain of the PM-localized PI4P-specific yeast phosphatase SAC1p can be used to specifically decrease the PI4P pool at the PM and therefore to determine the requirement of PI4P for the localization and function of proteins of interest^30,31,39^. We co-expressed the three AtADR1s (RNLs) and AtRPM1 (CNL) with SAC1 and determined their subcellular localization and protein abundance by confocal microscopy and western blot analysis, respectively. The N-terminally myristoylated and PM localized Arabidopsis CNL AtRPS5 was included as a control NLR as AtRPS5 PM localization (and function) was not expected to be affected by PM PI4P reduction^19,40^. Co-expression with the catalytically active wildtype SAC1 (SAC^WT^) affected the PM localization of all tested NLRs except AtRPS5 (Fig. 2a,c,e,g; Supplementary Fig. S2 a). By contrast, co-expression with the phosphatase dead mutant SAC1 (SAC1^dead^) protein had no influence on the PM localization of AtADR1, AtADR1-L1, AtADR1-L2, AtRPM1 and AtRPS5 (Fig. 2 a,c,e,g; Supplementary Fig. S2 a). Thus, the effect of SAC1 activity on the PM localization of AtRNLs and AtRPM1 is specific. Co-localization of AtADR1 with SAC1^WT^ at the PM was rarely detectable and the majority of AtADR1 was localized inside the cell, likely at the ER and/or cytosol (Fig. 2a). We also observed less AtADR1 accumulation in western blot analysis upon co-expression with SAC1^WT^ (Fig. 2b), indicating that most likely only the PM-localized pool of AtADR1 and not the ER-localized pool is affected by SAC1^WT^ co-expression. No fluorescence was observed for either AtADR1-L1 or AtRPM1, and only a very weak fluorescence for AtADR1-L2, after co-expression with SAC1^WT^ (Fig. 2c,e,g). This indicates that depleting the PM PI4P pool severely affects protein accumulation of AtADR1-L1, AtADR1-L2 and AtRPM1. Western blot analysis of AtADR1-L1, AtADR1-L2 and AtRPM1 upon co-expression with SAC1^WT^ confirmed the lack of NLR protein accumulation (Fig. 2d,f,h). A similar observation was previously reported for a phosphatidylserine specific binding protein, which is unstable in the Arabidopsis *pss1* mutant that lacks phosphatidylserine production^41^.

**Fig.2.**
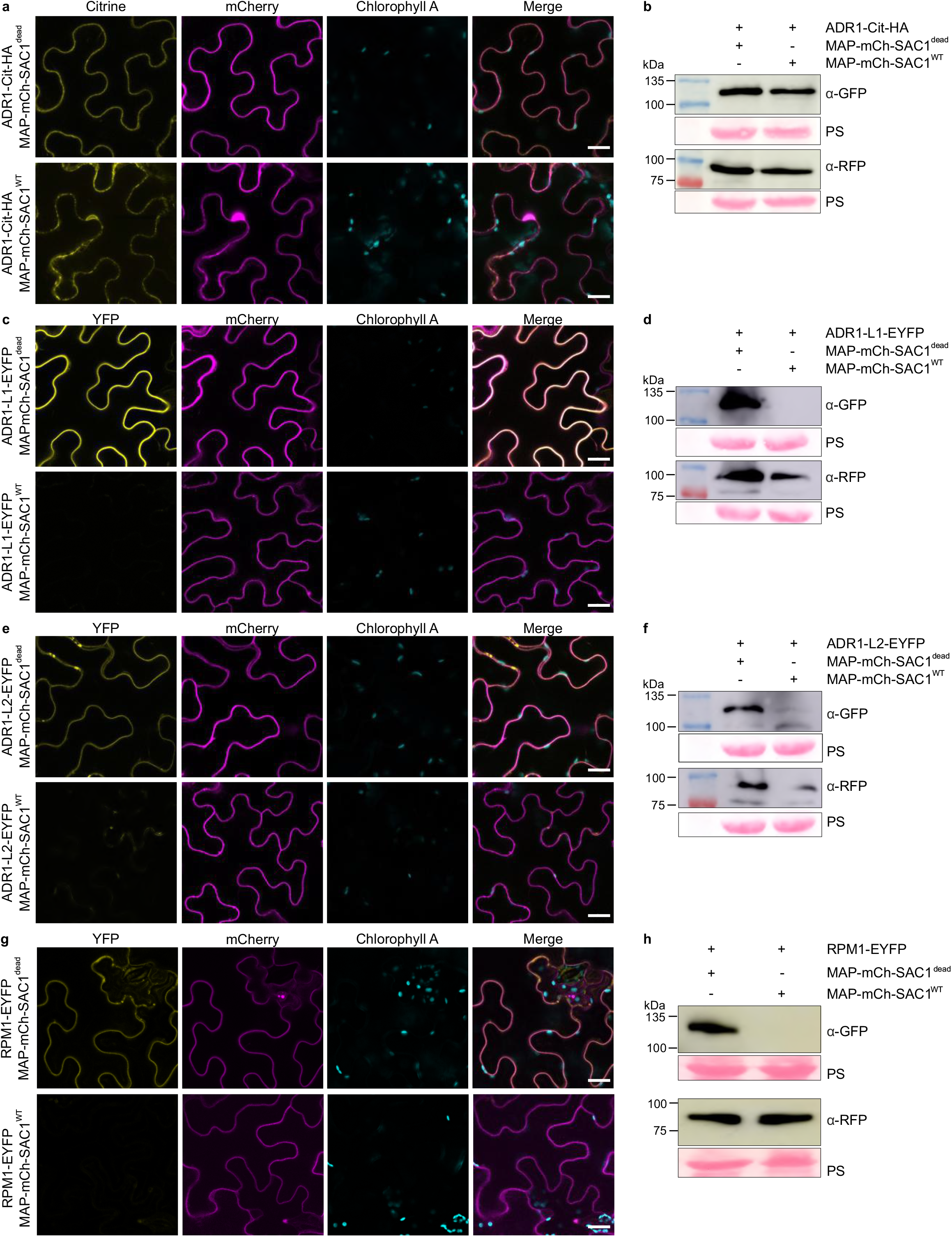
PI4P depletion reduces PM localization and stability of AtADR1s and AtRPM1. Effects on the localization and stability of AtADR1s (ADR1, ADR1-L1, ADR1-L2) and AtRPM1 after transient co-expression with SAC1^dead^ (upper panel) or SAC1^WT^ (lower panel). **a,** MAP-mCherry-SAC1^WT^ co-expression affects ADR1-Citrine-HA PM localization but not its endoplasmic reticulum localization. **c,e,g,** ADR1-L1-, ADR1-L2 and RPM1-EYFP fluorescence is not (c and g) or only weakly (e) detectable when co-expressed with MAP-mCherry-SAC1^WT^. Fusion proteins were transiently expressed in *N. benthamiana* leaves and confocal imaging was done at 4 hours after Estradiol induction (a), 2 days post infiltration (c, e) or 24 hours post infiltration (g). Localization of ADR1-Citrine-HA and ADR1-L1-, ADR1-L2 and RPM1-EYFP proteins is shown in the first column (Citrine/YFP, in yellow) and MAP-mCherry-SAC1^WT^ or MAP-mCherry-SAC1^dead^ is shown in the second column (mCherry, in magenta). Chloroplasts are shown in the third column (Chlorophyll A, in cyan) and the merged images are shown in the fourth column. Images are single plane secant views. Scale bars, 20 μm. **b,** Immunoblot analysis indicates a slightly reduced accumulation of ADR1-Citrine-HA after co-expression with MAP-mCherry-SAC1^WT^ compared to co-expression with MAP-mCherry-SAC1^dead^. **d,f,h,** Coexpression of ADR1-L1 (d), ADR1-L2 (f) and RPM1 (h) −EYFP with MAP-mCherry-SAC1^WT^ severely affects their stability. Immunoblot analysis of proteins infiltrated in (a, c, e, g) using anti-GFP and anti-RFP antibody are shown. Equal loading of the proteins is indicated by the Rubisco band from the Ponceau staining (PS). Protein samples were collected at 4 hours after Estradiol induction (b), 2 days post infiltration (d, f) or 24 hours post infiltration (h).

In order to test whether loss of NLR protein accumulation upon SAC1^WT^ co-expression was due to degradation of the mis-localized proteins we analysed protein levels by western blot in presence of protease and proteasome inhibitors. The specific inhibition of proteasomal degradation by Bortezomib (BTZ) had an observable effect on the accumulation of AtADR1-L2 (compare lane 2 with lane 6 in Supplementary Fig. S3c) and a weak effect on AtADR1 and AtADR1-L1 accumulation (Supplementary Fig. 3a,b). This indicates that proteasomal degradation is, at least partially, responsible for the degradation of mis-localized AtADR1s. In contrast, mis-localized AtRPM1 could not be stabilized in the presence of BTZ (Supplementary Fig. S3d), suggesting the proteasome plays no role in AtRPM1 degradation. This is consistent with previously published data^17^.

Together these results clearly demonstrate that all three AtADR1s and AtRPM1 require PI4P or a high electronegativity driven by PI4P at the PM for their proper localization and that loss of PM localization severely affects protein stability. Degradation of the mis-localized NLRs is, at least for the RNLs, partially mediated by the proteasome.

### Cell death function of Arabidopsis PM-localized RNLs and CNLs is PI4P dependent

PM localization of several NLRs, including AtRPM1, was shown to be important for their immune and cell death function^10,17–20^. The severe effect on the localization of the AtADR1s and AtRPM1 by depleting PI4P from the PM, prompted us to analyse whether their cell death function was also affected. Transient over-expression of the CC_R_ domains of the Arabidopsis RNLs ADR1, ADR1-L2 and NRG1.1 is sufficient to induce a cell death response in *N. benthamiana* (Fig. 3a-c)^16^. CC_R_ domain induced cell death activity was dramatically diminished by SAC1^WT^, but not SAC1^dead^ co-expression (Fig. 3a-c). These results suggest that the RNL CC_R_ domains induce cell death at the PM in a PI4P-dependent manner. Expression of the AtADR1-L1 CCR domain did not induce a visible cell death response in transient expression assays under our growth conditions and hence, could not be tested for PI4P dependency (Supplementary Fig. S4a). Interestingly, in contrast to the measurable negative effect of SAC1^WT^ activity on the accumulation of the full-length NLR proteins (Fig. 2b, d, f, h) we did not observe a similar effect on the CC_R_ domains (Fig. 3a-c). Altogether, PI4P depletion does not affect CC_R_ domain stability, but substantially affects CCR domain induced cell death.

**Fig.3.**
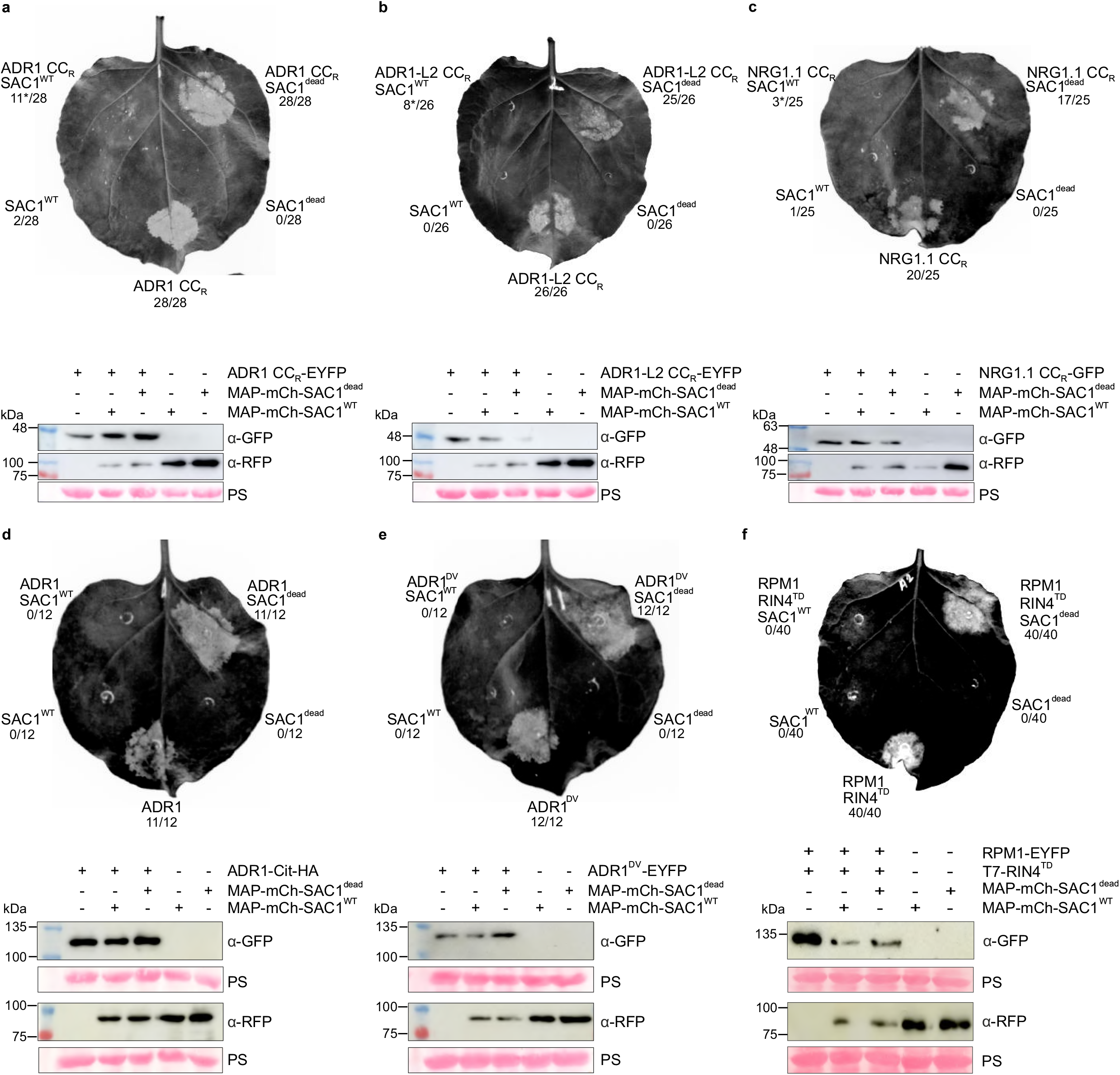
MAP-mCherry-SAC1 strongly affects AtADR1s and AtRPM1 cell death activity. Cell death activity of autoactive AtADR1s CC_R_ domains, full-length AtADR1 and the AtADR1^DV^ mutant as well as the phospho-mimic T7-RIN4^T166D^ activated RPM1 is suppressed by SAC1^WT^ co-expression. **a-f** (upper panels), Transient expression of ADR1 CC_R_ (a), ADR1-L2 CC_R_ (b), NRG1.1 CC_R_ (c), ADR1 (d), ADR1^D461V^ (e) and phospho-mimic T7-RIN4^T166D^ (RIN4^TD^)-activated RPM1 Citrine-HA- or EYFP-fusion proteins in *N. benthamiana* co-expressed with MAP-mCherry-SAC1^WT^ or MAP-mCherry-SAC1^dead^. Images of leaves were taken under UV light at 23 hours post infiltration (hpi) (a), 26 hpi (b), 28 hpi (c), 8 hp Estradiol induction (d), 30 hpi (e) and 24 hpi (f). Phospho-mimic T7-RIN4^TD^ was co-expressed to activate RPM1. White areas on the leaves indicate dead tissue. Numbers represent the number of leaves showing cell death out of the number of leaves analysed. Asterisk in (a and c) indicates weak cell death. **a-f** (lower panels), Immunoblot analysis of the proteins infiltrated in the upper panels using anti-GFP and anti-RFP (a-f) antibody. Membranes were horizontally cut into two pieces and probed with anti-GFP or anti-RFP antibody (a-c). Equal loading of the proteins is indicated by the Rubisco band from the Ponceau staining (PS). Protein samples were collected at 20 hpi (a-c), 24 hpi (d), 4 hp Estradiol induction (e) or 22 hpi (f).

Since the AtADR1 full-length protein induces a fast and strong cell death response in *N. benthamiana*, we further tested whether SAC1^WT^ co-expression can suppress cell death induced by full-length AtADR1 (Fig. 3 d). Similar to suppression of AtADR1 CC_R_ induced cell death by PI4P depletion, SAC1^WT^ expression also suppressed cell death activity of full-length AtADR1 (Fig. 3d).

To examine if suppression of cell death activity by SAC1^WT^ is not restricted to the CC_R_ domains and full-length wildtype AtADR1 we wanted to include the QHD motif mutants of the three AtADR1s in our cell death suppression experiments. However, only the AtADR1^DV^ mutant induced a strong, fast and reliable cell death response under our growth conditions (Fig. 1g). Consistent with the suppression of wildtype AtADR1 induced cell death by SAC1^WT^ co-expression, SAC1^WT^ co-expression also suppressed AtADR1^DV^ cell death activity (Fig. 3e). We conclude that PI4P depletion severely affects RNL and CC_R_ domain cell death activity, most likely due to loss of PM localization.

AtRPM1 guards the immune regulatory protein RIN4 (RPM1 INTERACTING PROTEIN 4) and is activated by an effector-triggered phosphorylation of RIN4 threonine 166^42,43^. AtRPM1 activation can be reconstituted in *N. benthamiana* by co-expression of AtRPM1 and a phosphomimic mutant of AtRIN4 (AtRIN4^T166D^)^17,44^. The strong cell death response upon AtRPM1 activation by AtRIN4^T166D^ was completely inhibited by SAC1^WT^ co-expression, but not by SAC1^dead^ (Fig. 3f). Cell death activity of effector-activated AtRPM1 was also severely affected by SAC1^WT^ co-expression (Supplementary Fig. S4b), demonstrating that AtRPM1 mediated cell death activity at the PM also depends on PI4P.

To demonstrate that the effect of decreasing the PM PI4P pool on cell death activity of the AtRNLs and AtRPM1 is specific and not a general effect on cell death induced by NLRs, we analysed whether SAC1^WT^ activity had any effect on cell death mediated by the myristoylated and “constitutively” PM localized AtRPS5. Similar to the AtRPM1 mediated cell death response, the AtRPS5 mediated and effector-triggered cell death can be reconstituted in transient expressions in *N. benthamiana*^45^. Neither the expression of SAC1^WT^ nor SAC1^dead^ suppressed effector-triggered and AtRPS5 mediated cell death (Supplementary Fig. S2e). These results suggest that the effect of SAC1 activity on cell death induction by the AtRNLs and AtRPM1 is specific.

Taken together, our results demonstrate that AtRNL and AtRPM1 cell death activity is significantly affected by PI4P depletion from the PM and further suggest that cell death activity of all AtRNLs, including the presumably ER-localized AtNRG1s^6,7^, takes place at the PM.

### PI4P depletion affects PM localization of AtADR1, AtADR1-L1 and AtADR1-L2 CC_R_ domains

Cell death activity of the AtADR1 and AtADR1-L2 CCR domain was notably diminished by SAC1^WT^ co-expression (Fig. 3a,b). However, unlike the full length AtADR1s, the stability of the AtADR1 CCR-domains was not affected (Fig. 3a,b; Supplementary Fig. S4a). To test whether PI4P depletion affects CC_R_ localization and hence function, we co-expressed the CC_R_ domains of all three AtADR1s with SAC1^WT^ or SAC1^dead^ and analysed their localization by confocal microscopy. All three CCR domains localized to the PM in the presence of SAC1^dead^ (Fig. 4; Supplementary Fig. S5). We also observed that the AtADR1 CC_R_ domains localized to dot-like structures and to ER membranes (Fig. 4a and Supplementary Fig. S5a). However, the PM localization of all three CCR domains was affected by SAC1^WT^ co-expression. Fluorescence of the CC_R_ domains was detected at intracellular puncta and also at ER membranes and/or the cytosol (Fig. 4; Supplementary Fig. S5). SAC1^WT^ triggered re-localization was more visible for AtADR1 and AtADR1-L2 CC_R_ domains than for the AtADR1-L1 CC_R_ domain (Supplementary Fig. S5).

**Fig.4.**
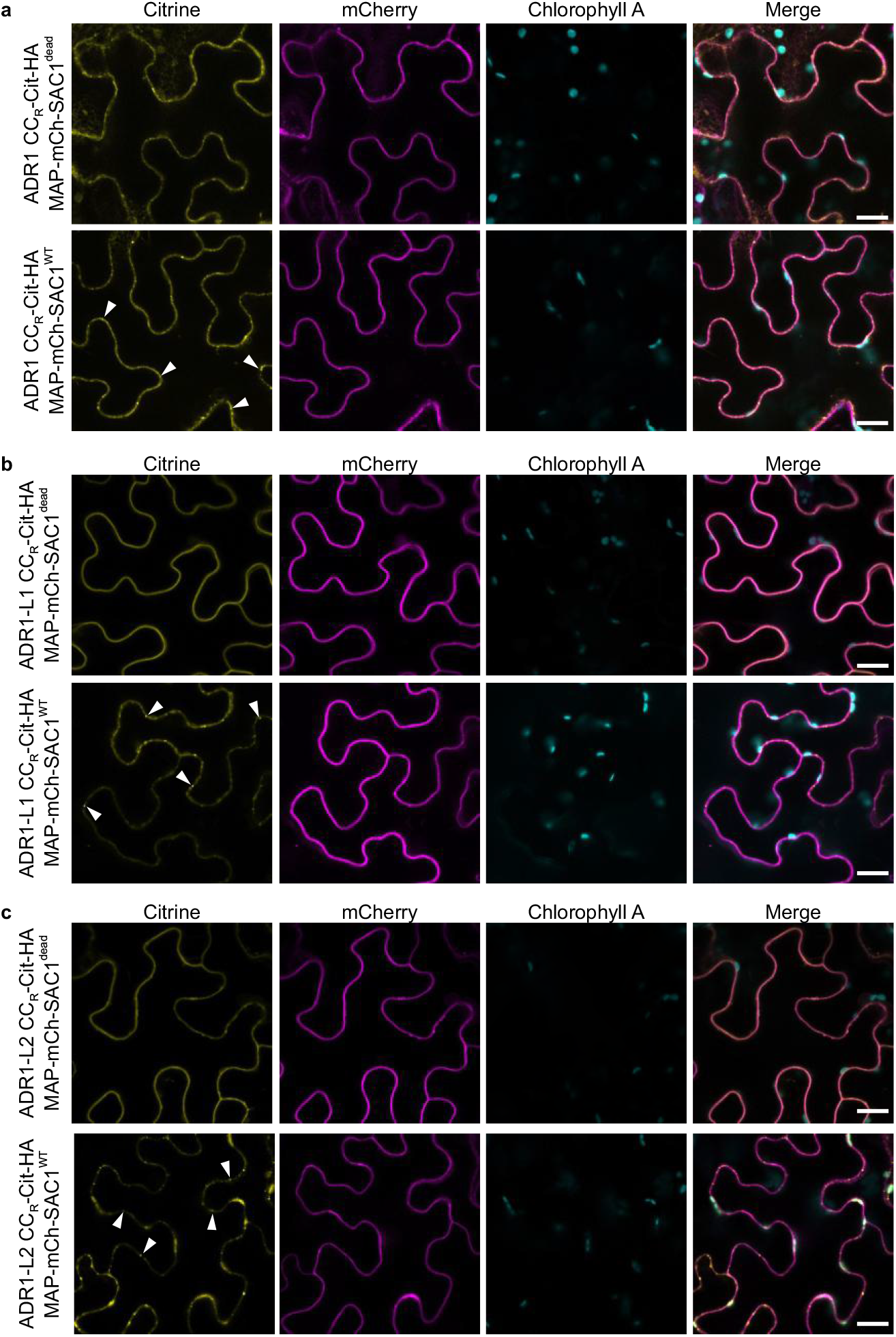
PI4P depletion affects PM localization of AtADR1 CC_R_ domains. Co-expression of SAC1^WT^ noticeably affects ADR1 CC_R_ (a), ADR1-L1 CC_R_ (b) and ADR1-L2 CC_R_ (c) localization. **a-c,** Citrine-HA tagged AtADR1 (ADR1, ADR1-L1, ADR1-L2) CC_R_ domains, MAP-mCherry-SAC1^dead^ (upper panels) or MAP-mCherry-SAC1^WT^ (lower panels) were transiently co-expressed in *N. benthamiana* leaves and confocal imaging was done at 3 hours (a) or 4 hours (b,c) post Estradiol induction. MAP-mCherry-SAC1^WT^ induces ADR1 CC_R_, ADR1-L1 CC_R_ and ADR1-L2 CC_R_ re-localization to intracellular puncta, most likely endosomes (white arrowheads in lower panels of a-c). Localization of ADR1 CCR-Cit-HA domains is shown with the first column (Citrine, in yellow) and MAP-mCherry-SAC1^dead^ or MAP-mCherry-SAC1^WT^ is shown in the second column (mCherry, in magenta). Chloroplasts are shown in the third column (Chlorophyll A, in cyan) and the merged images are shown in the fourth column (merge). Images are single plane secant views. Scale bars, 20 μm.

Thus, PI4P depletion from the PM leads to a reduced PM-localization (and loss of cell death function) of the CCR domains and potentially a (mis-)localization to endosomal compartments and the ER or cytosol. Proteins that are normally interacting with the PM in a PI4P- or electronegativity-dependent manner have been shown to ‘adopt’ endosomal localization once the PM PI4P pool is depleted^30,41^.

### AtADR1-, AtADR1-L1-, AtADR1-L2 CC_R_ and AtRPM1 CC domains specifically interact with anionic lipids *in vitro*

Reducing the abundance of PM PI4P levels negatively influenced the function, localization and stability of the tested Arabidopsis RNLs and RPM1. Thus, it is very likely that a direct interaction of AtADR1s and AtRPM1 with PM PI4P or other anionic lipids in general is required for their cell death activity. Given the structural homology of the CCR domain with the N-terminal HeLo domain of MLKL^21^ and the importance of the CC domain for cell death function of many CNLs^22^ we investigated whether the AtADR1s CCR and the AtRPM1 CC domains bind to specific phospholipids. We generated C-terminally haemagglutinin (HA)-tagged CC domain proteins *in vitro* and incubated the proteins on a lipid array (PIP strip). All three AtADR1 CC_R_ domains and the AtRPM1 CC domain directly interacted with PIPs, but not with other phospholipids or non-phosphorylated phosphoinositides (Fig. 5). A very weak interaction was also observed with the anionic and low abundant phosphatidylserine (PS) and phosphatidic acid (PA) (Fig. 5)^46^. These results suggest a strong binding of the AtRNLs and AtRPM1 CC_R_/CC domain with negatively charged PIPs most likely via an electrostatic interaction.

**Fig.5.**
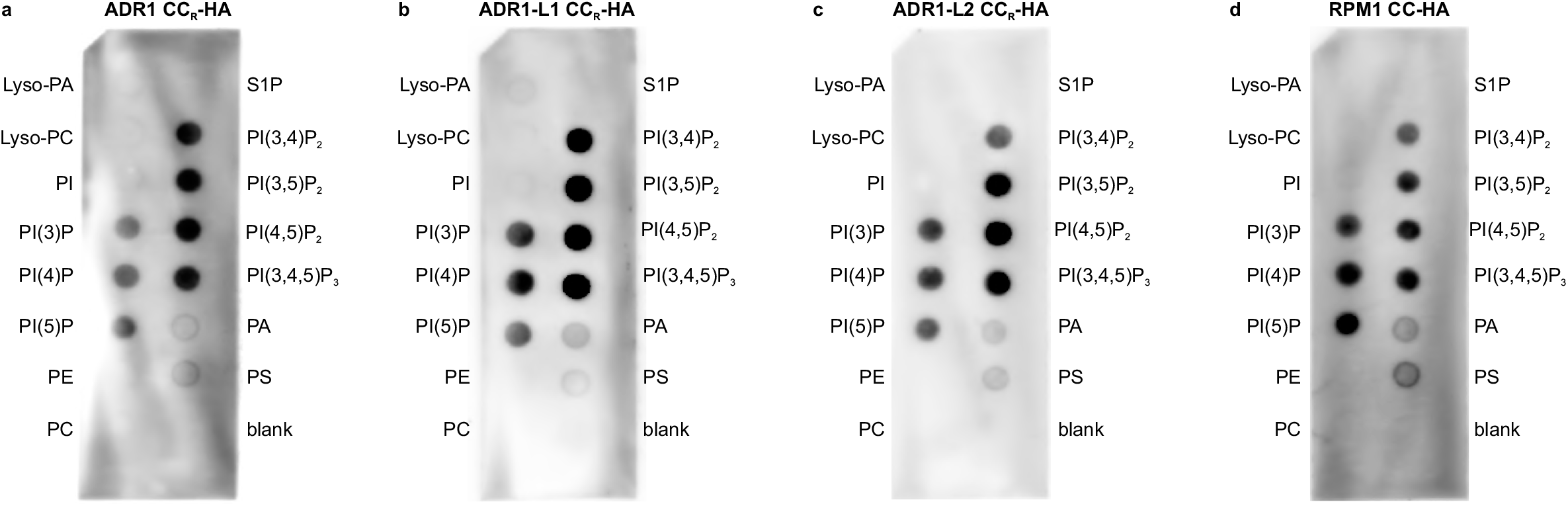
AtADR1s CC_R_ and AtRPM1 CC interact *in vitro* with anionic lipids. Arabidopsis ADR1s CCR and RPM1 CC domains can directly bind to anionic lipids *in vitro*. **a-d,** *In vitro* transcribed and translated AtADR1 (a), AtADR1-L1 (b) and AtADR1-L2 CC_R_ (c) and AtRPM1 CC (d) domains fused with a C-terminal single HA tag were incubated with a commercial PIP strip. Binding was analysed by immunoblotting with anti-HA antibody. The analysed CC domains bind strongly to PI(3)P, PI(4)P, PI(5)P, PI(3,4)P2, PI(3,5)P2, PI(4,5)P2 and PI(3,4,5)P3. A weak interaction was also detected with PA and PS.

### PI(4,5)P_2_ depletion has no impact on PM localization and cell death function of AtRNL and AtRPM1

The strong effect of PI4P depletion at the PM on the function and localization of AtRNLs and AtRPM1 and the specific interaction of their CC_R_/CC domain with anionic lipids (including PI4P) *in vitro*, suggest that PI4P plays a major role for their interaction with and function at the PM. However, phosphatidylinositol 4,5-bisphosphate (PI(4,5)P2) fulfils similar important cellular functions, is specifically found at the plant PM, and is also required for the interaction of many proteins with the PM^47^, like the mammalian MLKL proteins^26,27^. Although, PI(4,5)P2 is most likely not required for plant PM electronegativity^30^. Our observation of the additional direct binding of the AtADR1s CCR and AtRPM1 CC domains to PI(4,5)P2 (Fig. 5), prompted us to test whether PI(4,5)P2 is also required for AtRNL and AtCNL PM localization and cell death function. We co-expressed the PM-anchored wildtype PI(4,5)P2 5-phosphatase domain from the Drosophila OCRL protein (dOCRL^WT^) that specifically depletes the PI(4,5)P2 pool at the plant PM^47^ with AtADR1, AtADR1-L1, AtADR1-L2, AtRPM1 and AtRPS5. As a control we co-expressed a phosphatase dead mutant version of dOCRL (dOCRL^dead^) that is catalytically inactive^47^. Co-expression of neither dOCRL^WT^ nor dOCRL^dead^ had a visible effect on the (PM-) localization or protein expression of the tested AtRNLs and AtCNLs (Supplementary Figure S2c,d and S6). Co-expression of dOCRL^WT^ with the cell death inducing CC_R_ domains of AtADR1, AtADR1-L2 and AtNRG1.1 did not inhibit their activity and a strong cell death induction was visible for all three CC_R_ domains (Supplementary Fig. S7 a-c). Likewise, dOCRL^dead^ co-expression did not negatively affect the activity of the tested CC_R_ domains. Similarly, depleting the PI(4,5)P2 pool did not affect AtADR1 and AtADR1^DV^-induced cell death responses (Supplementary Fig. 7d,e). Further, we found no inhibition of the cell death activity of AtRPM1 or AtRPS5 in either the presence of dOCRL^WT^ or dOCRL^dead^ (Supplementary Fig. S4c, S2f and S7f). Consistent with the fact that PI(4,5)P2 depletion does not affect AtRNL and AtRPM1-mediated cell death, we also did not observe a negative effect on protein accumulation by PM PI(4,5)P2 depletion (Supplementary Fig. S6 and S7f). This suggests Arabidopsis RNLs, RPM1 and RPS5 cell death activity at the PM is independent of PI(4,5)P_2_.

Collectively, this demonstrates that PI(4,5)P2 is likely not a major contributor for AtADR1s (RNLs), AtRPM1 and AtRPS5 (CNL) localization and function at the PM.

## Discussion

The Arabidopsis CNL ZAR1 oligomerizes upon effector-induced activation, followed by a potential translocation to the PM where it is potentially forming a pore-like structure via the alpha 1 helix of its CC domain^10,48^. PM or endomembrane localization was shown to be necessary for the cell death and immune function of many CNLs^17,49^. Some CNLs localize to membranes via N-terminal myristoylation and/or palmitoylation, and the residues required for this post-translational modification were demonstrated to be important for CNL function^19^. However, the molecular mechanism underlying the localization of non-acylated PM/membrane-localized NLRs remains elusive. We present data that suggests a model in which AtRNLs and the CNL AtRPM1 require PI4P at the PM for proper localization, protein stability and cell death function upon (auto-) activation (Fig. 6). The localization is most likely regulated by direct binding of their CC/CC_R_ domain to anionic lipids (including the very abundant PI4P), possibly via positive charges in this domain. We cannot rule out the possibility that other mechanisms are also required, like the interaction with other (structural) lipids or proteins, e.g. integral membrane or transmembrane proteins. However, the strong effect of PI4P depletion from the PM on NLR function and stability, suggests that PI4P contributes significantly to RNL and CNL PM localization.

**Fig.6.**
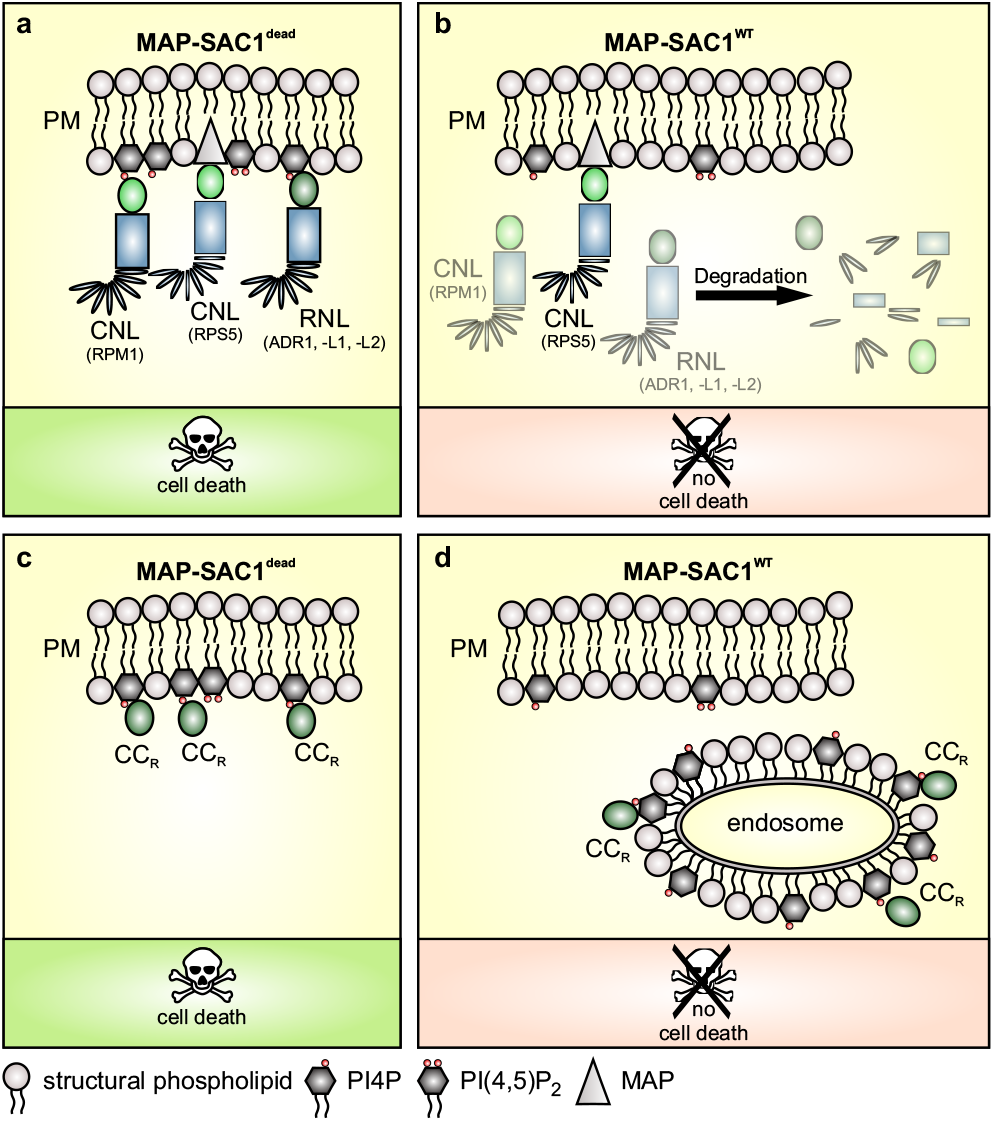
Proposed model of RNL and CNL localization and cell death/resistance function at the plasma membrane. Localization of RNLs and non-acylated CNLs, for example AtRPM1, to the plasma membrane (PM) is mediated by a direct interaction of their CCR or CC domains with anionic lipids, of which PI4P is the most abundant at the plant plasma membrane. **a,** Expression of catalytical inactive and forced PM-localized MAP-SAC1^DEAD^ does not affect RNL (ADR1s), myristoylated (RPS5) or non-acylated CNL (RPM1) PM localization, and consequently also not their cell death activity upon (auto-)activation. **b,** MAP-SAC1^WT^-mediated PI4P depletion from the PM severely affects RNL and non-acylated CNL, but not myristoylated CNL, localization. The decreased PI4P levels strongly affect PM electronegativity and this leads to a loss of binding to the PM and rapid degradation of RNLs and non-acylated CNLs. The reduced accumulation of RNLs and CNLs in the cell consequently leads to loss of RNL- and CNL-mediated cell death induction. **c,** The localization of RNL CC_R_ domains is not affected by MAP-SAC1^DEAD^ expression, similar to full-length RNLs. Thus, there is no observable effect on CCR domain autoactivity (cell death induction). **d,** PI4P depletion by MAP-SAC1^WT^ expression causes a relocalization of the CCR domains to intracellular puncta, probably endosomal compartments as their membranes might contain the highest electronegativity when MAP-SAC1^WT^ is expressed. This mis- or re-localization of CCR domains does not lead to their degradation. However, CCR cell death activity is severely reduced.

Interestingly, recent studies demonstrated that there is a reduction in PI4P and a specific enrichment of PI(4,5)P2 on interfacial membranes during successful infections, like the extra-haustorial membrane (EHM) in Arabidopsis powdery mildew infections, the extra-invasive hyphal membrane (EIHM) in Arabidopsis *Colletotrichum* infections or at the potato (*solanum tuberosum*) *Phytophthora infestans* infection sites^50–52^. The PI(4,5)P2 enrichment at the EHM and EIHM is an essential susceptibility factor, which is most likely pathogen-induced and requires the function of the host phosphatidylinositol 4-phosphate 5-kinases (PIP5K)^50,51^. It is possible that the depletion of PI4P and the simultaneous enrichment of PI(4,5)P2 at these host-pathogen interfaces result in a reduced accumulation of immune-regulatory proteins, for example NLRs, by removing possible binding sites and/or enhancing endocytosis of immune signaling components^50^. Plants however have evolved means to counteract this potentially pathogen/effector-induced enrichment of PI(4,5)P2 by downregulating the activity of PIP5Ks or upregulating the activity of phosphoinositide 5-phosphatases upon pathogen perception by cell-surface localized immune receptors^53,54^. Thus, actively changing or adjusting the lipid composition and homeostasis of the plant PM is part of the evolutionary arms race between the host and the pathogen. This indicates the importance of the regulation/manipulation of lipid homeostasis and the associated changes in protein localization/stability in this battle.

Likewise, a correlation between the lipid composition of the PM and immunity, as well as NLR (CNL) function and stability and an important function for phospholipase-dependent signalling in immunity was previously reported^55–59^. Plant phospholipase families C (PLCs) and D (PLDs) are involved in many aspects of abiotic and biotic stress responses^60^. However, the exact mechanisms of how these enzymes and their product(s) influence immunity are not well understood^61^. Perception of pathogen-derived danger signals by NLRs and cell-surface localized pathogen-recognition receptors (PRRs) lead to rapid recruitment and specific activation of PLDs and PLCs and to their recruitment to pathogen entry sites at the PM, as well as a biphasic transient Ca^2+^ influx^56,57,62^. PLDs and PLCs induce the production of inositol polyphosphates, phosphatidic acid (PA) and diacylglycerol (DAG), all of which can function as second messengers during immunity as well as other stress responses^61^. The PLC and PLD mediated generation of PA is required for NLR-triggered ROS production and HR, and external application of PA is sufficient to induce a cell death response and the transcriptional activation of the pathogen-responsive *PR1* promoter^58^. The hypothetical pore or ion (Ca^2+^)-channel forming capability of some CNLs at membranes is presumably required for their cell death activity and downstream immune signalling^21,63^. It is very likely that RNLs, having a RPW8-like/HeLo-like CC domain, use a similar mechanism for cell death induction and immunity. In light of our results it is tempting to hypothesize that (i) RNL (and most CNLs) activation leads to oligomerization and (enhanced or induced) interaction with PM/membrane anionic lipids, like PI4P, (ii) the formation of a transient Ca^2+^ channel/pore and the (iii) subsequent activation of calcium dependent and probably NLR-interacting phospholipases that in turn produce lipid messengers, such as PA and DAG, which (iv) might activate downstream signalling components required for cell death and resistance (Supplementary Figure 8)^10,21,55,58^.

## Methods

### Plasmid construction

The CDS from *ADR1* and *RPM1* were cloned into pENTR/D-TOPO (Thermo Fisher Scientific; Waltham, USA), while the CDS from *ADR1-L1, ADR1-L2, ADR1 CC_R_* (1-146aa)*, ADR1-L1 CC_R_* (1-155aa) *and ADR1-L2 CCR* (1-153aa) were cloned into pDONR221 (Invitrogen; Carlsbad, USA) generating pEntry clones by gateway cloning (Life Technologies; Carlsbad, USA). The corresponding point mutations for the QHV mutants *ADR1^D461V^, ADR1-L1^D489V^* and *ADR1-L2^D484V^* were introduced by site-directed mutagenesis PCR using primers listed in table S2. The PCR products were digested with *Dpn*I (NEB; Ipswich, USA) overnight and subsequently transformed into *Escherichia coli* DH5α. The CDS of RPS5 was cloned into pDONR207 (Invitrogen; Carlsbad, USA). The *NRG1.1 CC_R_* (1-180aa) CDS was synthesized with 3’ and 5’ gateway attachment sites into the pUC57Kan vector (Genescript, Piscataway NJ, USA). LR reactions (Gateway Cloning Technology, Life Technologies; Carlsbad, USA) were performed to introduce specific CDS into a modified Estradiol-inducible destination vector pMDC7-Citrine-HA^64^, the Estradiol-inducible destination vector pABindmCherry^65^ or the 35s-driven destination vector pGWB641^66^ as indicated. The 36 first amino acid of AtGPA1 (i.e. MAP sequence) were added to mCHERRYnoSTOP in pDONR207^67^ to generate MAP-mCHERRYnoSTOP in pDONR207. 2×35Sprom/pDONRP4-P1R^68^, MAP-mCHERRYnoSTOP in pDONR207 and SAC1in pDONR-P2R-P3 (or SAC1dead in pDONR-P2RP3)^30^ were recombined using LR reaction into pH7m34GW^69^ to generate 2×35Sprom::MAP-mCHERRY-SAC1 in pH7m34GW (or 2×35Sprom::MAP-mCHERRY-SAC1dead in pH7m34GW). 2×35Sprom in pDONRP4-P1R^68^, MAP-mCHERRYnoSTOP in pDONR207 and dOCRL in pDONR-P2R-P3 (or dOCRLdead in pDONR-P2RP3)^47^ were recombined using LR reaction into pH7m34GW^69^ to generate 2×35Sprom::MAP-mCHERRY-dOCRL in pH7m34GW (or 2×35Sprom::MAP-mCHERRY-dOCRLdead in pH7m34GW). Constructs were verified by sequencing and transformed into *Agrobacterium tumefaciens* strain GV3101 and used for transient expression in *Nicotiana benthamiana*.

### Transient expression in *N. benthamiana*

*Agrobacterium tumefaciens* strains were grown overnight at 28°C in LB media containing the appropriate antibiotics. The overnight cultures were centrifuged for 8 min at 8,500 rpm and the pellets were resuspended in induction buffer (10 mM MgCl2, 10 mM MES pH 5.6, 150 μM acetosyringone). The OD600 was adjusted to 0.05 (35S::P19) and 0.3 (35s::RPS5-EYFP, Dex::PBS1-3x-HA, Dex::AvrPphB-5x-myc, 35S::RPM1-EYFP, pRIN4::T7-RIN4^T166D^, Dex::AvrRpm1-HA, 35s::MAP-mCh-SAC1^WT/dead^, 35s::MAP-mCh-dOCRL^WT/dead^, 35s::ADR1/L1/L2-EYFP, Est::ADR1/L1/L2-Cit-HA, Est::ADR1 ^DV^/L1 ^DV^/L2^DV^-Cit-HA, Est::ADR1/L1 /L2-mCherry, Est::ADR1 ^DV^/L1 ^DV^/L2^DV^-mCherry, 35s::ADR1/L1/L2 CC, Est::ADR1/L1/L2 CC-Cit-HA) and samples were mixed as indicated. Agrobacteria mixtures were infiltrated into young leaves of 4-6 week old *N. benthamiana* WT plants using a 1-ml needleless syringe. The *N. benthamiana* plants were grown on soil under 12h light / 12h dark cycles (24°C/22°C, 70% humidity). Induction of protein expression was done 24 hours post infiltration using either 30 μM Dexamethasone (Sigma-Aldrich; St. Louis, USA) and 0.001% [v/v] Silwet L-77 or 20 μM Estradiol (Sigma-Aldrich; St. Louis, USA) and 0.001% [v/v] Silwet L-77 by spraying. Leaves were imaged for cell death or for protein localization at indicated timepoints.

### Chemical treatments

For PIC and BTZ treatments, *N. benthamiana* leaves were infiltrated with the indicated constructs using a 1 ml needleless syringe. At 23 hours post infiltration (hpi), leaves were infiltrated with induction buffer (10 mM MgCl2, 10 mM MES pH 5.6, 150 μM acetosyringone) only as Mock control or with induction buffer containing 2.5 μM BTZ (Santa Cruz Biotechnology; Dallas, USA) or 1x Halt™ Protease Inhibitor Cocktail (Thermo Fisher Scientific; Waltham, USA). For ADR1,20 μM Estradiol and 0.001% Silwet was infiltrated together with the Mock solution or the inhibitors to induce ADR1 expression. Leaf material was harvested 4 hours (ADR1) or 5 hours (ADR1-L1, ADR1-L2, RPM1) post inhibitor/mock treatment.

### HR/Cell Death Assay

Indicated constructs were transiently expressed in *N. benthamiana* leaves and leaves were imaged for cell death at the indicated time points. Cell death images were taken under UV light using the Amersham ImageQuant 800 western blot imaging system and an integrated Cy5 filter (GE Healthcare; Chalfont St. Giles, UK). Images were processed with Adobe Photoshop CS2 for adjustment of brightness and contrast. Note, since 35s::MAP-mCh-SAC1^WT^ often induces tissue collapse at around 52 hours post infiltration, cell death imaging has to be done at earlier timepoints.

### Confocal imaging

Protein localization was analysed at the indicated time points with the confocal laser scanning microscope LSM880 from Zeiss (Oberkochen, Germany), using a 40x or 63x water-immersion objective and the ZENblack software. EYFP and Citrine were excited using a 514 nm laser collecting emission between 516-556 nm; RFP and mCherry were excited using a 561 nm laser with an emission spectrum of 597-634 nm, Chlorophyll A was excited with a 561 nm laser and the emission spectrum was 661-682 nm. Focal plane images were processed with the ZENblue software (Zeiss) for adjustment of brightness and contrast. Maximum Z-projection images were processed with ImageJ.

### Western blot analysis of transiently expressed proteins

For protein extraction, 4 leaf discs (5mm diameter) were collected and frozen in liquid nitrogen, homogenized using a tissue homogenizer (Retsch GmbH) and resuspended in 190 μl grinding buffer (20 mM Tris-HCl pH 7, 150 mM NaCl, 1 mM EDTA pH 8, 1% [v/v] Triton X-100, 0.1% [w/v] SDS, 5 mM DTT, 1x Halt^™^ Protease Inhibitor Cocktail (Thermo Fisher Scientific; Waltham, USA)). Samples were incubated on ice for 5-10 min and then centrifuged for 15 min at 13,000 rpm and 4°C. 30 μl 5x SDS loading buffer (250 mM Tris-HCl pH 6.8, 50% [v/v] glycerol, 500 mM DTT, 10% [w/v] SDS, 0.005% [w/v] bromphenol blue) was added to 120 μl of supernatant. Proteins were denatured by incubation at 95°C for 5 min. Protein samples were resolved by electrophoresis on 8-10% SDS/PAGE gels, transferred to nitrocellulose membranes (GE Healthcare; Chalfont St Giles, UK) using semi-dry transfer (Bio-Rad Laboratories; Hercules, USA). Membranes were blocked in 5% [w/v] milk powder solved in 1x TBS with 1% [v/v] Tween-20 (TBS-T). Primary antibody incubations were done overnight at 4°C or for 1.5 hours at RT in 5% [w/v] milk powder diluted in TBS-T. Primary and secondary antibody dilutions were as follows: α-GFP 1:1500 (Roche Diagnostics; Basel, Switzerland), α-RFP 1:1000 (ChromoTek; Planegg-Martinsried, Germany), α-Myc 1:1000 (ChromoTek; Planegg-Martinsried, Germany), α-T7 Tag HRP conjugate 1:10.000 (Merck, Darmstadt, Germany), α-mouse HRP-conjugated 1:10.000 (Sigma-Aldrich; St. Louis, USA), α-rat HRP-conjugated 1:10.000 (Thermo Fisher Scientific; Waltham, USA). Chemiluminescence was detected using an Amersham Imager 600 or ImageQuant 800 (GE Healthcare; Chalfont St Giles, UK). Images were processed with Adobe Photoshop CS2 for adjustment of brightness and contrast.

### Co-immunoprecipitation

Frozen *N. benthamiana* leaf tissue (~200 mg) was collected and ground with a pre-cooled mortar and pestle with liquid nitrogen and then resuspended in 2.5 mL of extraction buffer (50mM HEPES pH 7.5, 50 mM NaCl, 10 mM EDTA pH 8.0, 0.5% [v/v] Triton X-100, 5 mM DTT, 1x Halt™ Protease Inhibitor Cocktail (Thermo Fisher Scientific; Waltham, USA)). Samples were kept for 10-30 min on ice and then cleared by centrifugation at 14,000 rpm for 5 min and 14,000 rpm for 15 min at 4°C. Protein extracts were incubated for 1 h with 25 μl GFP trap (ChromoTek; Planegg-Martinsried, Germany) on a rotating wheel at 4°C. Samples were captured by centrifugation at 2,400g at 4°C and washed two times with 1 ml of wash buffer (50 mM HEPES buffer pH 7.5, 150 mM NaCl, 10 mM EDTA pH 8.0, 0.2% [v/v] Triton X-100, 5 mM DTT, 1x Halt™ Protease Inhibitor Cocktail (Thermo Fisher Scientific; Waltham, USA)) by incubating the extracts for 5 min on a rotating wheel at 4°C and two additional times by inverting the tube six times. Bound proteins were eluted in 120 μl 2x SDS loading buffer (100 mM T ris-HCl pH 6.8, 20% [v/v] glycerol, 200 mM DTT, 4% [w/v] SDS, 0.002% [w/v] bromphenol blue) and denatured by boiling the proteins at 95°C for 5 min.

### *In vitro* transcription and translation and PIP strip assay

ADR1 CC_R_ (1-146aa), ADR1-L1 CC_R_ (1-155aa), ADR1-L2 CC_R_-HA (1-153aa) and RPM1 CC-HA (1-156aa) were expressed *in vitro* using the TnT^®^ SP6 High-Yield Wheat Germ Protein Expression System (Promega; Madison, USA) according to the manufacturer’s instructions. A PCR-generated DNA fragment was used as template for the transcription and translation reaction. Primers are listed in Table S2. Protein synthesis was confirmed on western blot using an HA-specific antibody. PIP strips (Echelon Biosciences; Salt Lake City, USA) were blocked overnight at 4°C in blocking buffer (PBS-T (0.1% [v/v] Tween-20), 4% [w/v] fatty acid-free BSA). 18 μl of the TnT reaction was added to 3 mL fresh blocking buffer and PIP strips were incubated for 1 h at RT with the protein. PIP strips were washed three times for 10 min with PBS-T. Binding of the proteins to the lipids was analysed by immunodetection using an HA-specific antibody. Primary antibody (α-HA 1:2000, Roche; Basel, Switzerland) incubation was done for 1 hour and 20 min at RT in blocking buffer, secondary antibody (α-rat HRP-conjugated 1:10.000; Thermo Fisher Scientific; Waltham, USA) incubation was done for 1 hour at RT in blocking buffer. Chemiluminescence was detected using the Amersham ImageQuant 800 (GE Healthcare; Chalfont St Giles, UK).

### Transmembrane and lipidation predictions

Full length protein sequences of Arabidopsis RPM1, RPS5, ADR1, ADR1-L1, ADR1-L2, NRG1.1 and NRG1.2 were used for prediction of potential transmembrane domains (TMDs) with three online tools: TMHMM2.0 (http://www.cbs.dtu.dk/services/TMHMM/)^70^, CCTOP (http://cctop.enzim.ttk.mta.hu/?_) and PredictProtein (https://predictprotein.org/)^71^, and for lipidation with the online tools: NBA-Palm (http://nbapalm.biocuckoo.org/)^72^, GPS-Palm (http://gpspalm.biocuckoo.cn/)^73^ and ExPASy Myristoylator (https://web.expasy.org/myristoylator/).

## Acknowledgments

We are grateful for technical support and experimental help from Christel Kulibaba-Mattern and Elke Sauberzweig. We thank Karin Schumacher for the VMA12 construct, Klaus Harter for the BRI1 construct and Roger Innes for PBS1-HA and AvrPphB-myc clones. We would also like to thank Thomas Stanislas, the El Kasmi lab, Nishimura lab and the Dangl Lab NLR group for critical comments and discussions on the project. We thank the University of Tübingen, the Deutsche Forschungsgemeinschaft (grant no. DFG-CRC1101 - project D09 to F.E.K.) and the Reinhard Frank Stiftung (Project ‘helperless plant’ to F.E.K.) for the financial support to F.E.K., the National Science Foundation (grant IOS-1758400 to M.T.N. and J.L.D.) and National Institutes of Health (grants GM107444 to J.L.D) for the financial support M.T.N. and J.L.D. M.T.N. was also supported by startup funds from Colorado State University, and J.L.D. is a Howard Hughes Medical Institute (HHMI) Investigator. Y.J. and M.C.C. were supported by ERC no. 3363360-APPL under FP/2007–2013 and ANR (caLIPSO; ANR-18-CE13-0025-02) to Y.J., ANR JC/JC Junior Investigator Grant (INTERPLAY; ANR-16-CE13-0021) and a SEED Fund ENS LYON-2016 to M.C.C.

## Author contributions

S.C.S. created RNL entry and destination constructs, performed confocal and cell death analysis for all RNLs, the in vitro transcription and translation assay, the PIP strip analysis and all western blot analysis for the RNL experiments and the BTZ treatments; F.M.A. performed confocal and cell death analysis for all RPM1 experiments; S.S. created RPS5 entry and destination constructs, performed cell death and western blot analysis for RPM1 and RPS5, and confocal analysis for RPS5.; A.B. did some cell death analysis for RNLs.; E.S., V.B. and L.W. assisted in creating RNL and CNL entry and destination constructs; Y.J. and M.C.C. provided unpublished SAC1 and dOCRL constructs; M.D. generated and characterized SAC1 and dOCRL constructs; S.C.S., F.M.A., M.T.N. and F.E.K conceived the study and designed the experiments; F.E.K. wrote the manuscript with help of S.C.S. and F.M.A.; S.S., A.B., Y.J., M.C.C., J.L.D. and M.T.N. reviewed and edited the manuscript.

## Additional information

Supplementary Table S1. Transmembrane domain and lipidation prediction summary for *Arabidopsis thaliana* RNLs and the CNL RPM1 and RPS5.

Supplementary Table S2. Primer list

## Supplementary Figure legends

**S1 Fig. AtADR1 proteins mainly localize to the PM.**

**a-h,** Maximum projection of Z-stack images clearly demonstrate AtADR1 proteins localization to the plasma membrane in transient expressions in *N. benthamiana* leaves. **a,b,e-h,** AtADR1 proteins (ADR1, ADR1-L1, ADR1-L2) localize mainly to the plasma membrane. The indicated ADR1 proteins fused to EYFP or Citrine-HA were transiently co-expressed with PM-resident BRI1-mRFP fusion-protein and confocal imaging was done at 4 hours (a, b, f) or 5 hours (h) post Estradiol induction or 2 days post infiltration (e, g). **c,d.** ADR1 also localizes to the endoplasmic reticulum (ER). Wildtype ADR1 or ADR1^DV^ fused to Citrine-HA was transiently co-expressed with the ER-localized VMA12-RFP fusion-protein and confocal imaging was done at 3 hours (c) and 4 hours (d) post Estradiol induction. Localization of EYFP and Citrine-HA tagged ADR1 proteins is shown with the first column (Citrine/YFP, in yellow) and the co-localized PM-localized BRI1-mRFP or ER-localized VMA12-RFP is shown in the second column (RFP, in magenta). Chloroplasts are shown in the third column (Chlorophyll A, in cyan) and the merged images are shown in the fourth column. Images shown here are a maximum projection of Z-stack images. Scale bars, 20 μm.

**S2 Fig. RPS5 localization and cell death activity at the plasma membrane is not affected by MAP-SAC1 or MAP-dOCRL co-expression.**

**a,** Plasma membrane localization of RPS5-EYFP is not affected by co-expression of MAP-mCherry-SAC1^dead^ (upper panel) or MAP-mCherry-SAC1^WT^ (lower panel). **c,** Co-expression of MAP-mCherry-dOCRL^dead^ (upper panel) or MAP-mCherry-dOCRL^WT^ (lower panel) does not affect RPS5-EYFP PM localization. Indicated proteins were transiently expressed in *N. benthamiana* leaves and confocal imaging was done at 24 hours post infiltration. Localization of RPS5-EYFP proteins is shown with the first column (YFP, in yellow) and MAP-mCherry-SAC1^WT^, MAP-mCherry-SAC1^dead^, MAP-mCherry-dOCRL^dead^ and MAP-mCherry-dOCRL^WT^ are shown in the second column (mCherry, in magenta). Chloroplasts are shown in the third column (Chlorophyll A, in cyan) and the merged images are shown in the fourth column (merge). Images are single plane secant views. Scale bars, 20 μm. **b,d,** Immunoblot analysis of the proteins infiltrated in (a) and (c) using anti-GFP and anti-RFP antibody. Equal loading of the proteins is indicated by the Rubisco band from the Ponceau staining (PS). Samples were collected 24 hours post infiltration. **e,f,** Effector (AvrPphB)-triggered and RPS5-EYFP mediated cell death is not suppressed by MAP-mCherry-SAC1^WT^ (e) or MAP-mCherry-dOCRL^WT^ (f) co-expression in *N. benthamiana*. (e) and (f) upper panels, leaf images showing cell death induction of activated RPS5-EYFP. Transient expression of Dexamethasone-inducible AvrPphB-MYC and PBS1-HA with constitutively expressed RPS5-EYFP, MAP-mCherry-SAC1^WT^ or MAP-mCherry-SAC1^dead^ (e) or MAP-mCherry-dOCRL^dead^ and MAP-mCherry-dOCRL^WT^ (f). Leaf images were taken under UV light at 24 hours post Dexamethasone induction, which corresponds to 2 days post infiltration (e,f). AvrPphB-MYC and PBS1-HA expression was induced with 30 μM Dexamethasone to activate RPS5-EYFP. White areas indicate dead leaf tissue. Numbers represent the number of leaves showing cell death out of the number of leaves analysed. (e) and (f) lower panels, Immunoblot analysis of the proteins infiltrated in the upper panels using anti-GFP and anti-RFP antibody. Equal loading of the proteins is indicated by the Rubisco band from the Ponceau staining (PS). Protein samples were collected at 6 hours post Dexamethasone induction, which corresponds to 28 hours post infiltration.

**S3 Fig. Degradation of mis-localized AtRPM1 and AtADR1 proteins is not or only partially blocked by proteasome inhibitors.**

**a-c,** Bortezomib (BTZ) treatment partially inhibits degradation of mis-localized ADR1 proteins. **d,** Mislocalized RPM1 protein degradation can neither be blocked by a protease inhibitor cocktail (PIC) nor BTZ. Shown are immunoblot analysis of ADR1 (a), ADR1-L1 (b), ADR1-L2 (c) and RPM1 (d) Citrine-HA or EYFP fusion proteins that were transiently co-expressed with MAP-mCherry-SAC1^WT^ or MAP-mCherry-SAC1^dead^ in *N. benthamiana* using anti-GFP antibody. Equal loading of the proteins is indicated by the Rubisco band from the Ponceau staining (PS). Samples were collected 4 hours (a) or 5 hours (b-d) post inhibitor and Estradiol (a) treatments, which corresponds to 27 (a) and 25 hours (b-d) post infiltration.

**S4 Fig. Effector-triggered AtRPM1-mediated cell death is reduced by PI4P depletion**

**a** upper panel, CC_R_ domain of AtADR1-L1 induces no visible cell death symptoms and thus no effect of MAP-mCherry-SAC1^WT^ co-expression on AtADR1-L1 CCR activity is observable. Transient co-expression of ADR1-L1 CCR-EYFP, MAP-mCherry-SAC1^WT^ or MAP-mCherry-SAC1^DEAD^ in *N. benthamiana* leaves. **b** upper panel, mCherry-SAC1^WT^ co-expression noticeably reduced cell death activity of AvrRpm1-HA activated RPM1-EYFP. Transient expression of RPM1-EYFP, AvrRPM1-HA and MAP-mCherry-SAC1^WT^ or MAP-mCherry-SAC1^DEAD^ in *N. benthamiana* leaves. **c** upper panel, Cell death activity of AvrRpm1-HA activated RPM1-EYFP was not blocked by co-expression of MAP-mCherry-dOCRL^dead^ or MAP-mCherry-dOCRL^WT^. Transient expression of RPM1-EYFP, AvrRPM1-HA and MAP-mCherry-dOCRL^dead^ or MAP-mCherry-dOCRL^WT^ in *N. benthamiana* leaves. AvrRPM1-HA expression was induced with 30 μM Dexamethasone 20 hours post infiltration. Leaf images were taken under UV light at 24 hours post infiltration (a) or 24 hours post Dexamethasone induction (b and c). White areas indicate dead leave tissue. Numbers represent the number of leaves showing cell death out of the number of leaves analysed. Asterisk in (b) indicates weak cell death. **a-c** lower panels, Immunoblot analysis of the proteins infiltrated in leaves shown in upper panels using anti-GFP and anti-RFP antibody. Equal loading of the proteins is indicated by the Rubisco band from the Ponceau staining (PS). Protein samples were collected at 20 hours post infiltration (a) and 6 hours post Dexamethasone induction, which corresponds to 26 hours post infiltration (b and c).

**S5 Fig. PI4P depletion affects the PM localization of Arabidopsis ADR1 CC_R_ domains**

**a-c,** Co-expression SAC1^WT^ affects ADR1 CC_R_ (a), ADR1-L1 CC_R_ (b) and ADR1-L2 CC_R_ (c) localization. Citrine-HA tagged AtADR1 (ADR1, ADR1-L1, ADR1-L2) CC_R_ domains were transiently co-expressed with MAP-mCherry-SAC1^dead^ (a-c upper panel) or MAP-mCherry-SAC1^WT^ (a-c lower panel) in *N. benthamiana* leaves. Confocal imaging was done at 3 hours (a) or 4 hours after Estradiol induction (b,c). ADR1 CC_R_, ADR1-L1 CC_R_ and ADR1-L2 CC_R_ domains re-localization to intracellular puncta, potentially endosomes, is indicated (white arrow heads). Localization of ADR1 CCR-Citrine-HA domains is shown with the first column (Citrine, in yellow) and MAP-mCherry-SAC1^WT^ or MAP-mCherry-SAC1^dead^ is shown in the second column (mCherry, in magenta). Chloroplasts are shown in the third column (Chlorophyll A, in cyan) and the merged images are shown in the fourth column (merge). Images shown here are a maximum projection of Z-stack images. Scale bars, 20 μm.

**S6 Fig. PI(4,5)P_2_ is not required for the PM localization and stability of AtADR1s and AtRPM1.**

**a, c, e, g,** Plasmam membrane localization of AtADR1s (ADR1-Citrine-HA, ADR1-L1-EYFP, ADR1-L2-EYFP) and AtRPM1-EYFP is not altered when dOCRL^dead^ (upper panel) or dOCRL^WT^ (lower panel) is co-expressed. Proteins were transiently expressed in *N. benthamiana* leaves and confocal imaging was done 3 hours post Estradiol induction (a), 2 days post infiltration (c, e) or 24 hours post infiltration (g). Localization of Citrine-HA/-EYFP tagged ADR1s and RPM1-EYFP is shown with the first column (Citrine/YFP, in yellow) and MAP-mCherry-dOCRL^WT^ or MAP-mCherry-dOCRL^dead^ is shown in the second column (mCherry, in magenta). Chloroplasts are shown in the third column (Chlorophyll A, in cyan) and the merged images are shown in the fourth column (merge). Images are single plane secant views. Scale bars, 20 μm. **b, d, f, h,** Immunoblot analysis of proteins infiltrated in (a, c, e, g) using anti-GFP and anti-RFP antibody show no affect on NLR stability by dOCRL^WT^ or dOCRL^dead^ co-expression. Equal loading of proteins is indicated by the Rubisco band from the Ponceau staining (PS). Samples were collected at 4 hours post Estradiol induction (b), 2 days post infiltration (d, f) or 24 hours post infiltration (h).

**S7 Fig. AtADR1s and AtRPM1 cell death activity is not affected by depletion of PI(4,5)P2 from the plasma membrane via MAP-dOCRL co-expression.**

**a-f,** dOCRL^WT^ co-expression does not affect the cell death induced by ADR1 CC_R_ (a), ADR1-L2 CC_R_ (b), NRG1.1 CC_R_ (c) domains, full-length ADR1 (d), mutant ADR1^DV^ (e) or RPM1 (f). **a-f upper panels**, Transient expression of Citrine-HA or EYFP tagged autoactive, ADR1 (d), ADR1^D461V^ mutant (e) and phospho-mimic T7-RIN4^T166D^ activated RPM1 (f) co-expressed with MAP-mCherry-dOCRLWT or MAP-mCherry-dOCRL^dead^ in *N. benthamiana*. Leaf images were taken under UV light at 24 hours post infiltration (hpi) (a), 26 hpi (b), 28 hpi (c), 9 hours post Estradiol induction (d), 30 hpi (e) and 24 hpi (f). Phospho-mimic T7-RIN4^T166D^ (RIN4^TD^) was co-expressed to activate RPM1. White areas in leaves indicate dead tissue. Numbers represent the number of leaves showing cell death out of the number of leaves analysed. Asterisk in (c) indicates weak HR. **a-f lower panels,** Immunoblot analysis of proteins infiltrated in the upper panels using anti-GFP and anti-RFP antibody. Equal loading of proteins is indicated by the Rubisco band from the Ponceau staining (PS). Samples were collected at 20 hpi (a-c), 24 hpi (d), 4 hours post Estradiol induction (e) or 22 hpi (f).

**S8 Fig. Proposed model of AtRNL localization, oligomerization and function during immunity.**

Arabidopsis RNLs constitutively localize at the plasma membrane through the interaction of their CCR domains with anionic lipids, including PI4P. (1) RNL activation, either by pathogen infection or autoactivating mutations, leads to conformational changes inducing oligomerization, (2) the formation of a transient Ca^2+^ channel/pore and the (3) subsequent recruitment or activation of calcium dependent and probably NLR-interacting phospholipases that (4) in turn produce lipid messengers, such as PA and DAG, which (5) might activate downstream signalling components required for NLR-mediated (6) cell death and resistance outputs. The lipase-like protein EDS1 (ENHANCED DISEASE SUSCEPTIBLE 1) and its sequence-related direct partners SAG101 (SENESCENCE ASSOCIATED GENE 101) and PAD4 (PHYTOALEXIN DEFICIENT 4) are key immune regulators of NLR-mediated immunity, but also of basal resistance.

**Supplementary Table 1.**
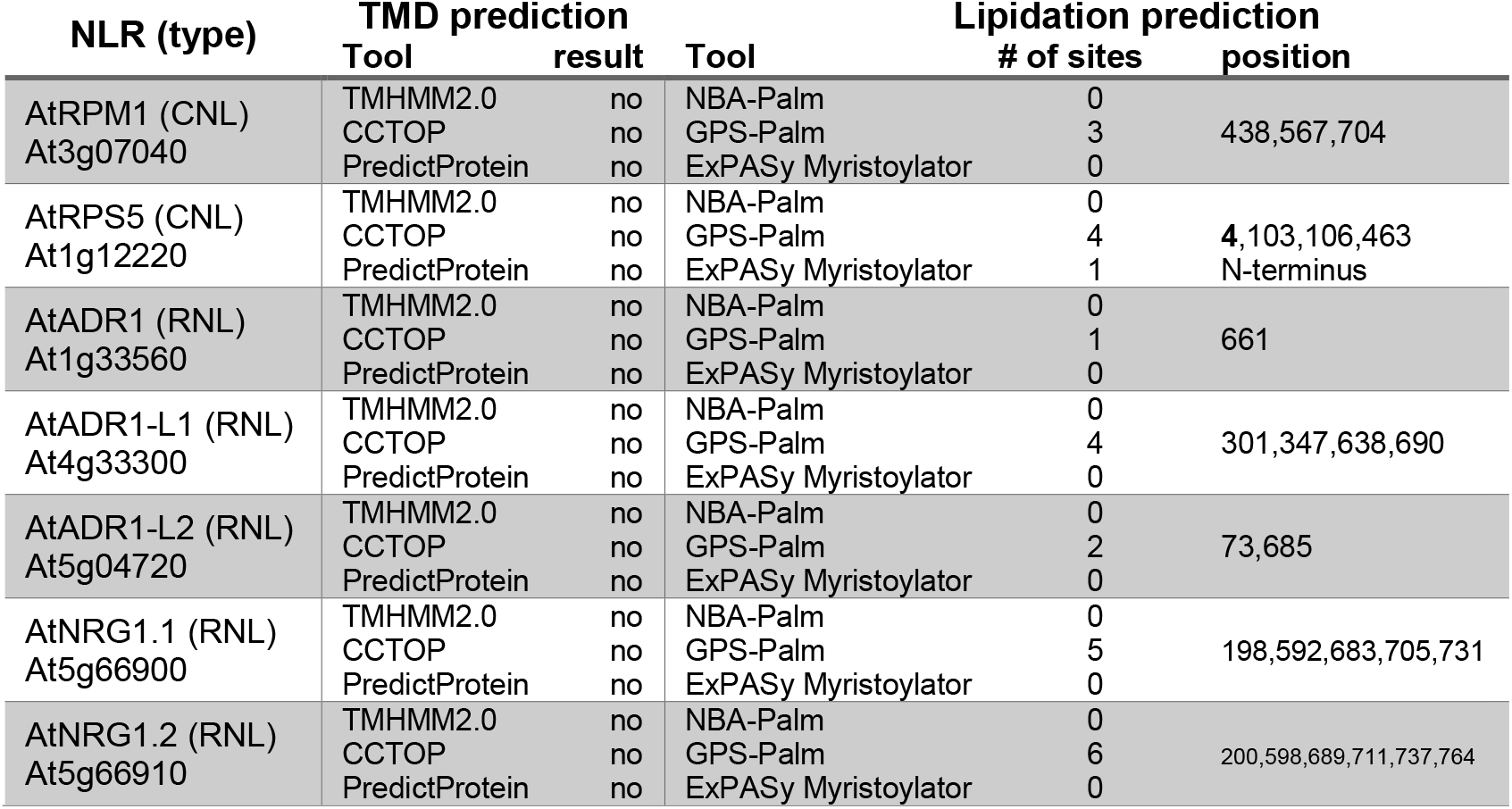
Transmembrane domain and lipidation prediction summary for *Arabidopsis thaliana* RNLs and the CNL RPM1 and RPS5.

**Supplementary Table 2.**
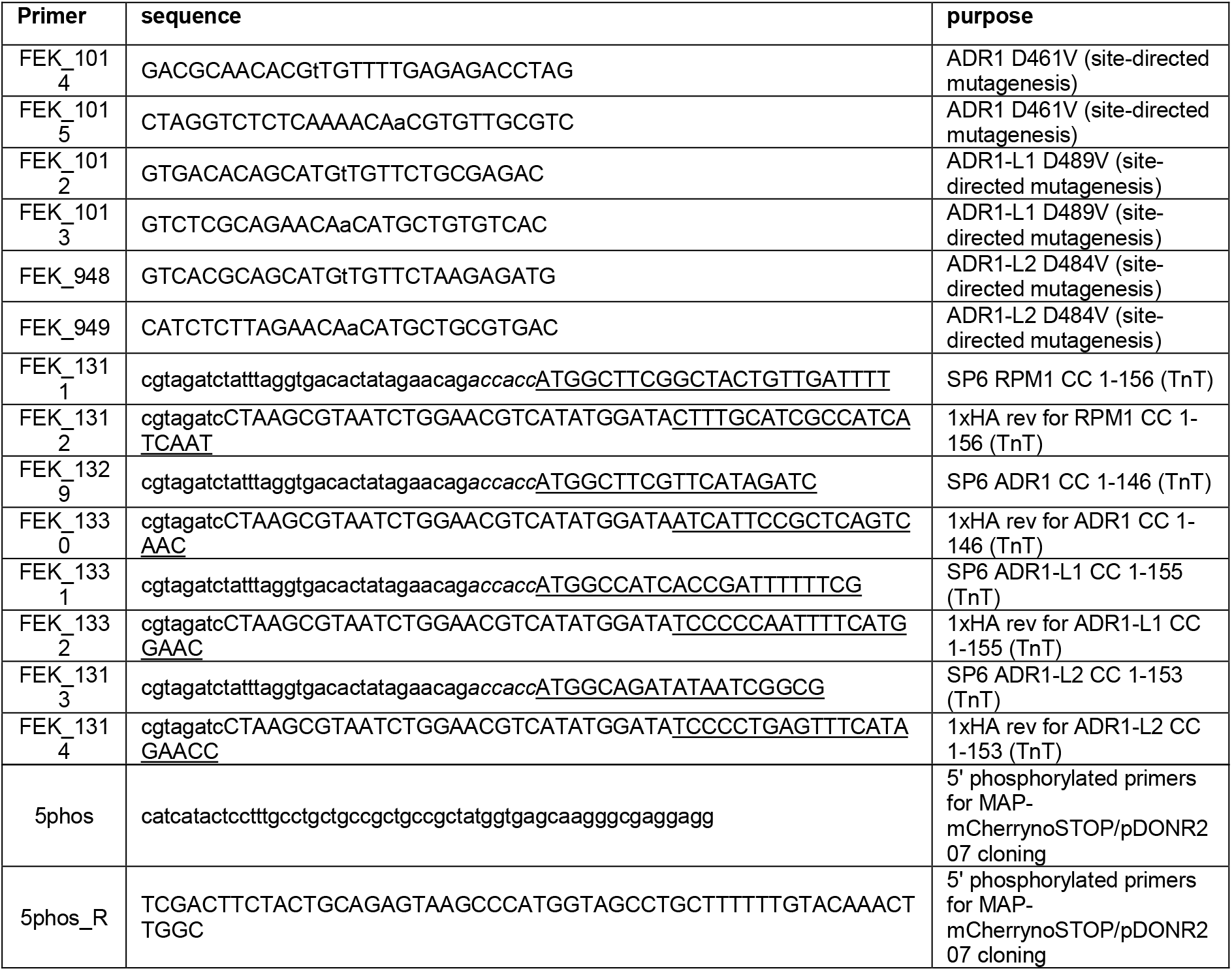
Primer list.

## References

1 Monteiro, F. & Nishimura, M. T. Structural, Functional, and Genomic Diversity of Plant NLR Proteins: An Evolved Resource for Rational Engineering of Plant Immunity. Annu Rev Phytopathol 56, 243–267, doi:10.1146/annurev-phyto-080417-045817 (2018).

2 Jones, J. D. & Dangl, J. L. The plant immune system. Nature 444, 323–329, doi:10.1038/nature05286 (2006).

3 Balint-Kurti, P. The plant hypersensitive response: concepts, control and consequences. Mol Plant Pathol 20, 1163–1178, doi:10.1111/mpp.12821 (2019).

4 Bonardi, V. et al. Expanded functions for a family of plant intracellular immune receptors beyond specific recognition of pathogen effectors. Proc Natl Acad Sci U S A 108, 16463–16468, doi:10.1073/pnas.1113726108 (2011).

5 Castel, B. et al. Diverse NLR immune receptors activate defence via the RPW 8-NLR NRG 1. New Phytologist (2018).

6 Wu, Z. et al. Differential regulation of TNL-mediated immune signaling by redundant helper CNL s. New Phytologist (2018).

7 Lapin, D. et al. A Coevolved EDS1-SAG101-NRG1 Module Mediates Cell Death Signaling by TIR-Domain Immune Receptors. Plant Cell 31, 2430–2455, doi:10.1105/tpc.19.00118 (2019).

8 Qi, T. et al. NRG1 functions downstream of EDS1 to regulate TIR-NLR-mediated plant immunity in Nicotiana benthamiana. Proceedings of the National Academy of Sciences 115, E10979–E10987, doi:10.1073/pnas.1814856115 (2018).

9 Saile, S. C. et al. Two unequally redundant “helper” immune receptor families mediate Arabidopsis thaliana intracellular “sensor” immune receptor functions. PLoS Biol 18, e3000783, doi:10.1371/journal.pbio.3000783 (2020).

10 Wang, J. et al. Reconstitution and structure of a plant NLR resistosome conferring immunity. Science 364, doi:10.1126/science.aav5870 (2019).

11 Hu, M., Qi, J., Bi, G. & Zhou, J. M. Bacterial Effectors Induce Oligomerization of Immune Receptor ZAR1 In Vivo. Mol Plant 13, 793–801, doi:10.1016/j.molp.2020.03.004 (2020).

12 Li, L., Habring, A., Wang, K. & Weigel, D. Atypical Resistance Protein RPW8/HR Triggers Oligomerization of the NLR Immune Receptor RPP7 and Autoimmunity. Cell Host Microbe 27, 405–417 e406, doi:10.1016/j.chom.2020.01.012 (2020).

13 Martin, R. et al. Structure of the activated Roq1 resistosome directly recognizing the pathogen effector XopQ. bioRxiv, 2020.2008.2013.246413, doi:10.1101/2020.08.13.246413 (2020).

14 Burdett, H. et al. The Plant “Resistosome”: Structural Insights into Immune Signaling. Cell Host Microbe 26, 193–201, doi:10.1016/j.chom.2019.07.020 (2019).

15 Xiong, Y., Han, Z. & Chai, J. Resistosome and inflammasome: platforms mediating innate immunity. Curr Opin Plant Biol 56, 47–55, doi:10.1016/j.pbi.2020.03.010 (2020).

16 Collier, S. M., Hamel, L. P. & Moffett, P. Cell death mediated by the N-terminal domains of a unique and highly conserved class of NB-LRR protein. Mol Plant Microbe Interact 24, 918–931, doi:10.1094/MPMI-03-11-0050 (2011).

17 Gao, Z., Chung, E. H., Eitas, T. K. & Dangl, J. L. Plant intracellular innate immune receptor Resistance to Pseudomonas syringae pv. maculicola 1 (RPM1) is activated at, and functions on, the plasma membrane. Proc Natl Acad Sci U S A 108, 7619–7624, doi:10.1073/pnas.1104410108 (2011).

18 El Kasmi, F. et al. Signaling from the plasma-membrane localized plant immune receptor RPM1 requires self-association of the full-length protein. Proc Natl Acad Sci U S A 114, E7385–E7394, doi:10.1073/pnas.1708288114 (2017).

19 Qi, D., DeYoung, B. J. & Innes, R. W. Structure-function analysis of the coiled-coil and leucine-rich repeat domains of the RPS5 disease resistance protein. Plant Physiol 158, 1819–1832, doi:10.1104/pp.112.194035 (2012).

20 Wang, J. et al. Plant NLR immune receptor Tm-22 activation requires NB-ARC domain-mediated self-association of CC domain. PLoS Pathog 16, e1008475, doi:10.1371/journal.ppat.1008475 (2020).

21 Jubic, L. M., Saile, S., Furzer, O. J., El Kasmi, F. & Dangl, J. L. Help wanted: helper NLRs and plant immune responses. Curr Opin Plant Biol 50, 82–94, doi:10.1016/j.pbi.2019.03.013 (2019).

22 Bentham, A. R., Zdrzałek, R., De la Concepcion, J. C. & Banfield, M. J. Uncoiling CNLs: Structure/function approaches to understanding CC domain function in plant NLRs. Plant and Cell Physiology (2018).

23 Daskalov, A. et al. Identification of a novel cell death-inducing domain reveals that fungal amyloid-controlled programmed cell death is related to necroptosis. Proc Natl Acad Sci U S A 113, 2720–2725, doi:10.1073/pnas.1522361113 (2016).

24 Hofmann, K. The Evolutionary Origins of Programmed Cell Death Signaling. Cold Spring Harb Perspect Biol, doi:10.1101/cshperspect.a036442 (2019).

25 Murphy, J. M. The Killer Pseudokinase Mixed Lineage Kinase Domain-Like Protein (MLKL). Cold Spring Harb Perspect Biol 12, doi:10.1101/cshperspect.a036376 (2020).

26 Quarato, G. et al. Sequential Engagement of Distinct MLKL Phosphatidylinositol-Binding Sites Executes Necroptosis. Mol Cell 61, 589–601, doi:10.1016/j.molcel.2016.01.011 (2016).

27 Dondelinger, Y. et al. MLKL compromises plasma membrane integrity by binding to phosphatidylinositol phosphates. Cell Rep 7, 971–981, doi:10.1016/j.celrep.2014.04.026 (2014).

28 Heo, W. D. et al. PI(3,4,5)P3 and PI(4,5)P2 lipids target proteins with polybasic clusters to the plasma membrane. Science 314, 1458–1461, doi:10.1126/science.1134389 (2006).

29 McLaughlin, S. & Murray, D. Plasma membrane phosphoinositide organization by protein electrostatics. Nature 438, 605–611, doi:10.1038/nature04398 (2005).

30 Simon, M. L. et al. A PtdIns(4)P-driven electrostatic field controls cell membrane identity and signalling in plants. Nat Plants 2, 16089, doi:10.1038/nplants.2016.89 (2016).

31 Gronnier, J. et al. Structural basis for plant plasma membrane protein dynamics and organization into functional nanodomains. Elife 6, doi:10.7554/eLife.26404 (2017).

32 Friedrichsen, D. M., Joazeiro, C. A., Li, J., Hunter, T. & Chory, J. Brassinosteroid-insensitive-1 is a ubiquitously expressed leucine-rich repeat receptor serine/threonine kinase. Plant Physiol 123, 1247–1256, doi:10.1104/pp.123.4.1247 (2000).

33 Viotti, C. et al. The endoplasmic reticulum is the main membrane source for biogenesis of the lytic vacuole in Arabidopsis. Plant Cell 25, 3434–3449, doi:10.1105/tpc.113.114827 (2013).

34 Williams, S. J. et al. An autoactive mutant of the M flax rust resistance protein has a preference for binding ATP, whereas wild-type M protein binds ADP. Mol Plant Microbe Interact 24, 897–906, doi:10.1094/MPMI-03-11-0052 (2011).

35 Van Ooijen, G. et al. Structure-function analysis of the NB-ARC domain of plant disease resistance proteins. Journal of Experimental Botany 59, 1383–1397, doi:10.1093/jxb/ern045 (2008).

36 Roberts, M., Tang, S., Stallmann, A., Dangl, J. L. & Bonardi, V. Genetic requirements for signaling from an autoactive plant NB-LRR intracellular innate immune receptor. PLoS Genet 9, e1003465, doi:10.1371/journal.pgen.1003465 (2013).

37 Wang, J. & Chai, J. Molecular actions of NLR immune receptors in plants and animals. Sci China Life Sci 63, 1–14, doi:10.1007/s11427-019-1687-6 (2020).

38 Boyes, D. C., Nam, J. & Dangl, J. L. The Arabidopsis thaliana RPM1 disease resistance gene product is a peripheral plasma membrane protein that is degraded coincident with the hypersensitive response. Proc Natl Acad Sci U S A 95, 15849–15854, doi:10.1073/pnas.95.26.15849 (1998).

39 Doumane, M. & Caillaud, M. C. Assessing Extrinsic Membrane Protein Dependency to PI4P Using a Plasma Membrane to Endosome Relocalization Transient Assay in Nicotiana benthamiana. Methods Mol Biol 2177, 95–108, doi:10.1007/978-1-0716-0767-1_9 (2020).

40 Pottinger, S. E. & Innes, R. W. RPS5-Mediated Disease Resistance: Fundamental Insights and Translational Applications. Annu Rev Phytopathol 58, 139–160, doi:10.1146/annurev-phyto-010820-012733 (2020).

41 Platre, M. P. et al. A Combinatorial Lipid Code Shapes the Electrostatic Landscape of Plant Endomembranes. Dev Cell 45, 465–480 e411, doi:10.1016/j.devcel.2018.04.011 (2018).

42 Chung, E. H. et al. Specific threonine phosphorylation of a host target by two unrelated type III effectors activates a host innate immune receptor in plants. Cell Host Microbe 9, 125–136, doi:10.1016/j.chom.2011.01.009 (2011).

43 Liu, J., Elmore, J. M., Lin, Z. J. & Coaker, G. A receptor-like cytoplasmic kinase phosphorylates the host target RIN4, leading to the activation of a plant innate immune receptor. Cell Host Microbe 9, 137–146, doi:10.1016/j.chom.2011.01.010 (2011).

44 Chung, E. H., El-Kasmi, F., He, Y., Loehr, A. & Dangl, J. L. A plant phosphoswitch platform repeatedly targeted by type III effector proteins regulates the output of both tiers of plant immune receptors. Cell Host Microbe 16, 484–494, doi:10.1016/j.chom.2014.09.004 (2014).

45 Ade, J., DeYoung, B. J., Golstein, C. & Innes, R. W. Indirect activation of a plant nucleotide binding site-leucine-rich repeat protein by a bacterial protease. Proc Natl Acad Sci U S A 104, 2531–2536, doi:10.1073/pnas.0608779104 (2007).

46 Colin, L. A. & Jaillais, Y. Phospholipids across scales: lipid patterns and plant development. Curr Opin Plant Biol 53, 1–9, doi:10.1016/j.pbi.2019.08.007 (2020).

47 Doumane, M. et al. iDePP: a genetically encoded system for the inducible depletion of PI(4,5)P<sub>2</sub> in <em>Arabidopsis thaliana</em>. bioRxiv, 2020.2005.2013.091470, doi:10.1101/2020.05.13.091470 (2020).

48 Wang, J. et al. Ligand-triggered allosteric ADP release primes a plant NLR complex. Science 364, doi:10.1126/science.aav5868 (2019).

49 Engelhardt, S. et al. Relocalization of late blight resistance protein R3a to endosomal compartments is associated with effector recognition and required for the immune response. Plant Cell 24, 5142–5158, doi:10.1105/tpc.112.104992 (2012).

50 Qin, L. et al. Specific Recruitment of Phosphoinositide Species to the Plant-Pathogen Interfacial Membrane Underlies Arabidopsis Susceptibility to Fungal Infection. Plant Cell 32, 1665–1688, doi:10.1105/tpc.19.00970 (2020).

51 Shimada, T. L. et al. Enrichment of Phosphatidylinositol 4,5-Bisphosphate in the Extra-Invasive Hyphal Membrane Promotes Colletotrichum Infection of Arabidopsis thaliana. Plant Cell Physiol 60, 1514–1524, doi:10.1093/pcp/pcz058 (2019).

52 Rausche, J. et al. A phosphoinositide 5-phosphatase from Solanum tuberosum is activated by PAMP-treatment and may antagonize phosphatidylinositol 4,5-bisphosphate at Phytophthora infestans infection sites. New Phytol, doi:10.1111/nph.16853 (2020).

53 Menzel, W. et al. A PAMP-triggered MAPK cascade inhibits phosphatidylinositol 4,5-bisphosphate production by PIP5K6 in Arabidopsis thaliana. New Phytol 224, 833–847, doi:10.1111/nph.16069 (2019).

54 Addad, F. et al. Management of patients with acute ST-elevation myocardial infarction: Results of the FAST-MI Tunisia Registry. PLoS One 14, e0207979, doi:10.1371/journal.pone.0207979 (2019).

55 Yuan, X., Wang, Z., Huang, J., Xuan, H. & Gao, Z. Phospholipidase Ddelta Negatively Regulates the Function of Resistance to Pseudomonas syringae pv. Maculicola 1 (RPM1). Front Plant Sci 9, 1991, doi:10.3389/fpls.2018.01991 (2018).

56 Schloffel, M. A. et al. The BIR2/BIR3-Associated Phospholipase Dgamma1 Negatively Regulates Plant Immunity. Plant Physiol 183, 371–384, doi:10.1104/pp.19.01292 (2020).

57 Johansson, O. N. et al. Redundancy among phospholipase D isoforms in resistance triggered by recognition of the Pseudomonas syringae effector AvrRpm1 in Arabidopsis thaliana. Front Plant Sci 5, 639, doi:10.3389/fpls.2014.00639 (2014).

58 Andersson, M. X., Kourtchenko, O., Dangl, J. L., Mackey, D. & Ellerstrom, M. Phospholipase-dependent signalling during the AvrRpm1-and AvrRpt2-induced disease resistance responses in Arabidopsis thaliana. Plant J 47, 947–959, doi:10.1111/j.1365-313X.2006.02844.x (2006).

59 Bargmann, B. O. & Munnik, T. The role of phospholipase D in plant stress responses. Curr Opin Plant Biol 9, 515–522, doi:10.1016/j.pbi.2006.07.011 (2006).

60 Hong, Y. et al. Plant phospholipases D and C and their diverse functions in stress responses. Prog Lipid Res 62, 55–74, doi:10.1016/j.plipres.2016.01.002 (2016).

61 Li, J. & Wang, X. Phospholipase D and phosphatidic acid in plant immunity. Plant Sci 279, 45–50, doi:10.1016/j.plantsci.2018.05.021 (2019).

62 Xing, J. et al. Secretion of Phospholipase Ddelta Functions as a Regulatory Mechanism in Plant Innate Immunity. Plant Cell 31, 3015–3032, doi:10.1105/tpc.19.00534 (2019).

63 Wang, J., Chern, M. & Chen, X. Structural dynamics of a plant NLR resistosome: transition from autoinhibition to activation. Sci China Life Sci 63, 617–619, doi:10.1007/s11427-019-9536-x (2020).

64 Curtis, M. D. & Grossniklaus, U. A gateway cloning vector set for high-throughput functional analysis of genes in planta. Plant Physiol 133, 462–469, doi:10.1104/pp.103.027979 (2003).

65 Bleckmann, A., Weidtkamp-Peters, S., Seidel, C. A. & Simon, R. Stem cell signaling in Arabidopsis requires CRN to localize CLV2 to the plasma membrane. Plant Physiol 152, 166–176, doi:10.1104/pp.109.149930 (2010).

66 Nakamura, S. et al. Gateway binary vectors with the bialaphos resistance gene, bar, as a selection marker for plant transformation. Biosci Biotechnol Biochem 74, 1315–1319, doi:10.1271/bbb.100184 (2010).

67 Jaillais, Y., Belkhadir, Y., Balsemao-Pires, E., Dangl, J. L. & Chory, J. Extracellular leucine-rich repeats as a platform for receptor/coreceptor complex formation. Proc Natl Acad Sci U S A 108, 8503–8507, doi:10.1073/pnas.1103556108 (2011).

68 Marques-Bueno, M. D. M. et al. A versatile Multisite Gateway-compatible promoter and transgenic line collection for cell type-specific functional genomics in Arabidopsis. Plant J 85, 320–333, doi:10.1111/tpj.13099 (2016).

69 Karimi, M., Bleys, A., Vanderhaeghen, R. & Hilson, P. Building blocks for plant gene assembly. Plant Physiol 145, 1183–1191, doi:10.1104/pp.107.110411 (2007).

70 Sonnhammer, E. L., von Heijne, G. & Krogh, A. A hidden Markov model for predicting transmembrane helices in protein sequences. Proc Int Conf Intell Syst Mol Biol 6, 175–182 (1998).

71 Yachdav, G. et al. PredictProtein--an open resource for online prediction of protein structural and functional features. Nucleic Acids Res 42, W337–343, doi:10.1093/nar/gku366 (2014).

72 Xue, Y., Chen, H., Jin, C., Sun, Z. & Yao, X. NBA-Palm: prediction of palmitoylation site implemented in Naive Bayes algorithm. BMC Bioinformatics 7, 458, doi:10.1186/1471-2105-7-458 (2006).

73 Ning, W. et al. GPS-Palm: a deep learning-based graphic presentation system for the prediction of S-palmitoylation sites in proteins. Brief Bioinform, doi:10.1093/bib/bbaa038 (2020).

