## Supplementary figures and images for "Coiled-coil and RPW8-type immune receptors function at the plasma membrane in a phospholipid dependent manner"

### Supplementary Figures S1-S8

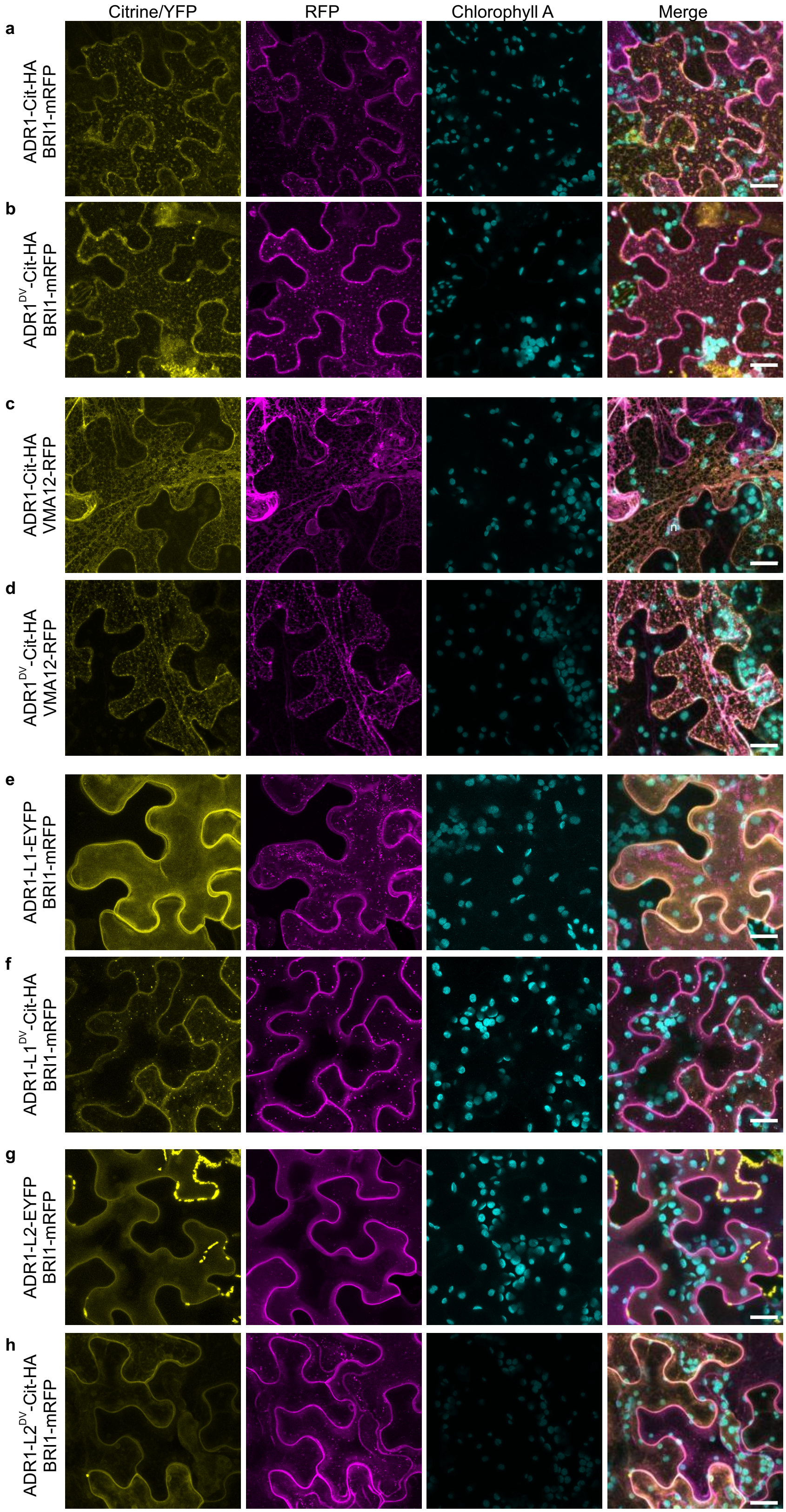

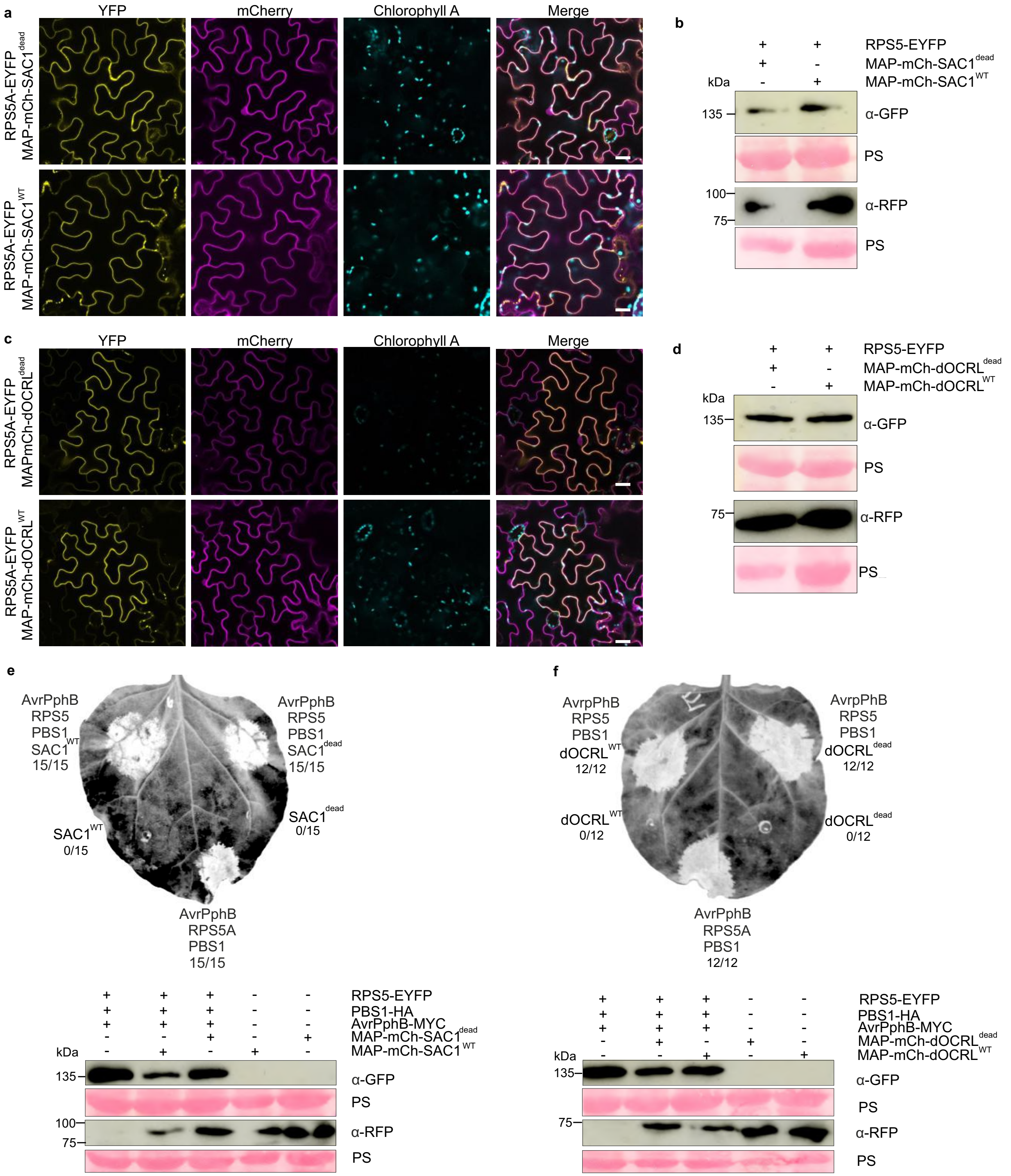

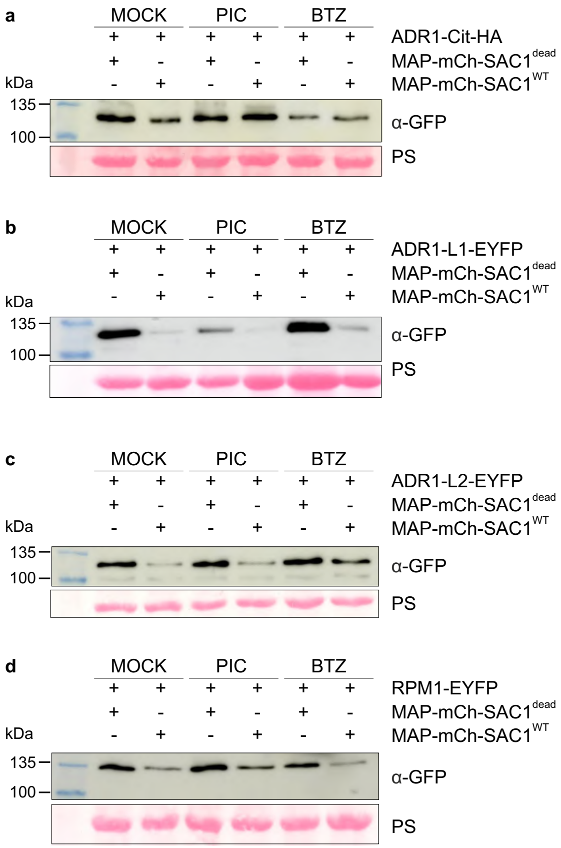

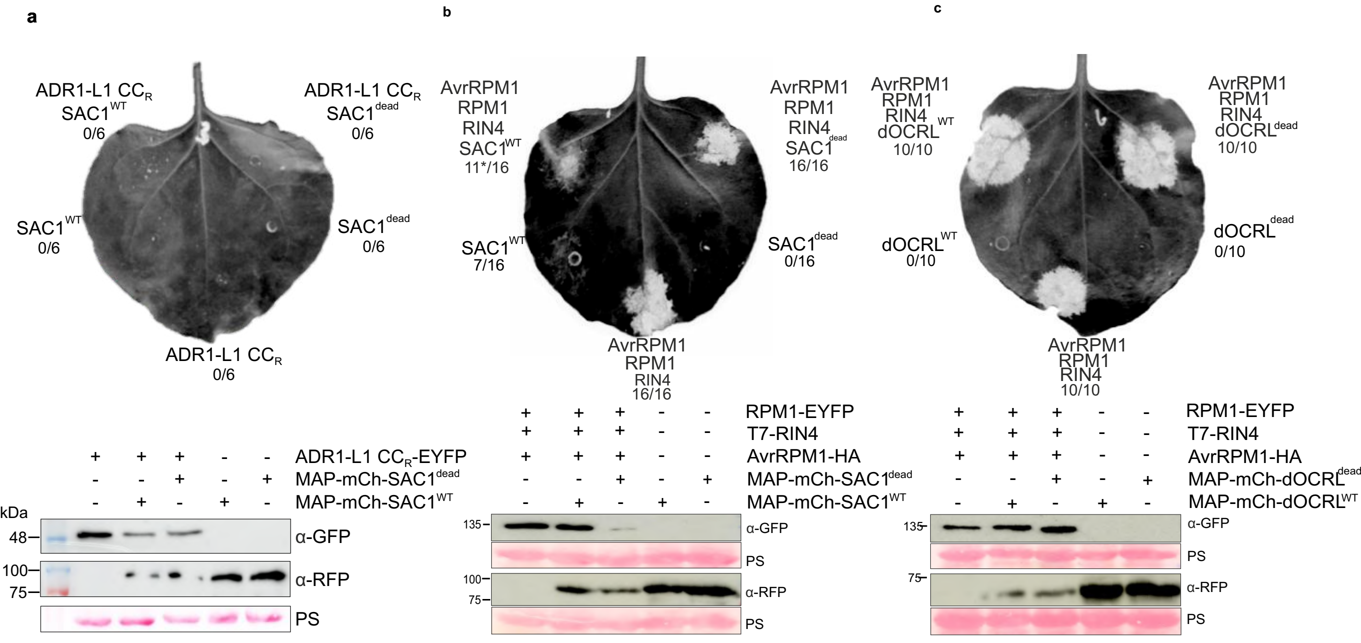

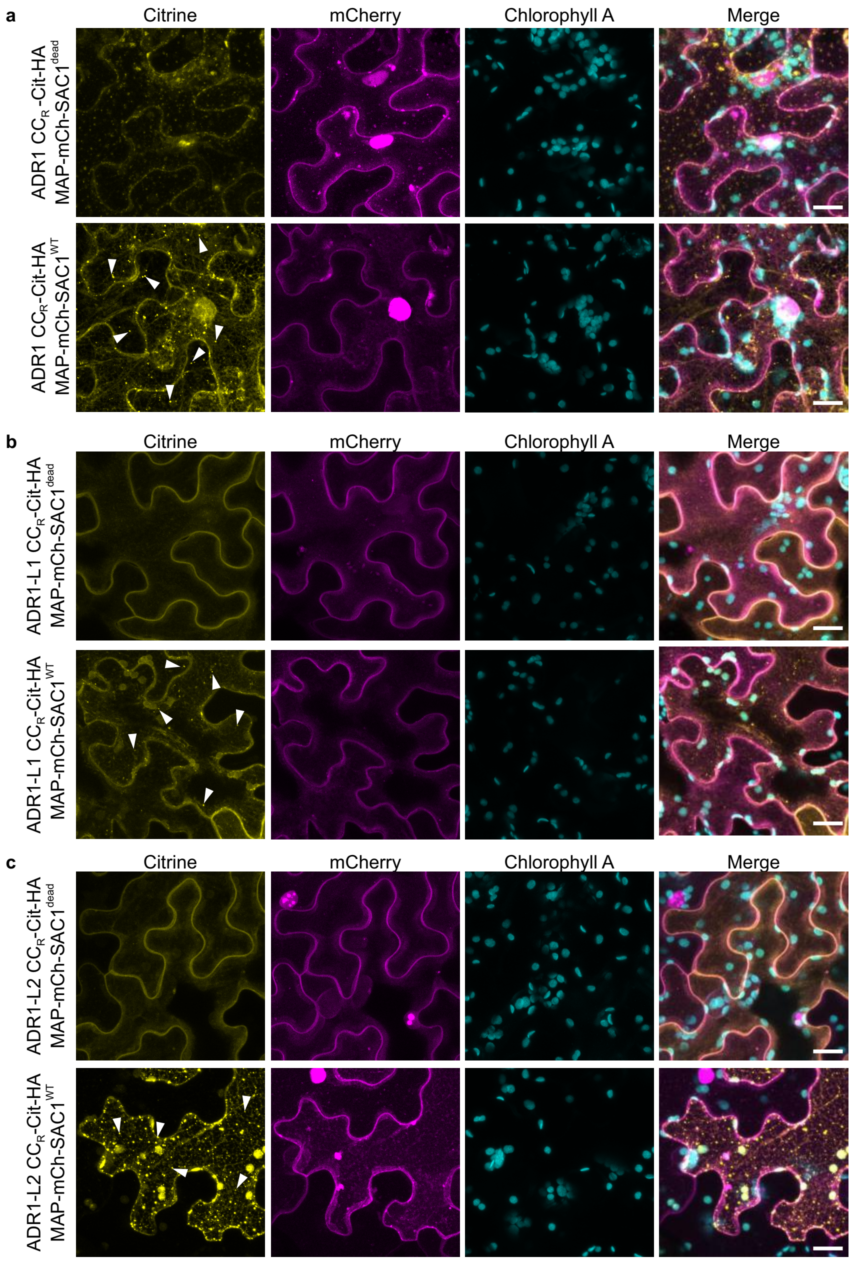

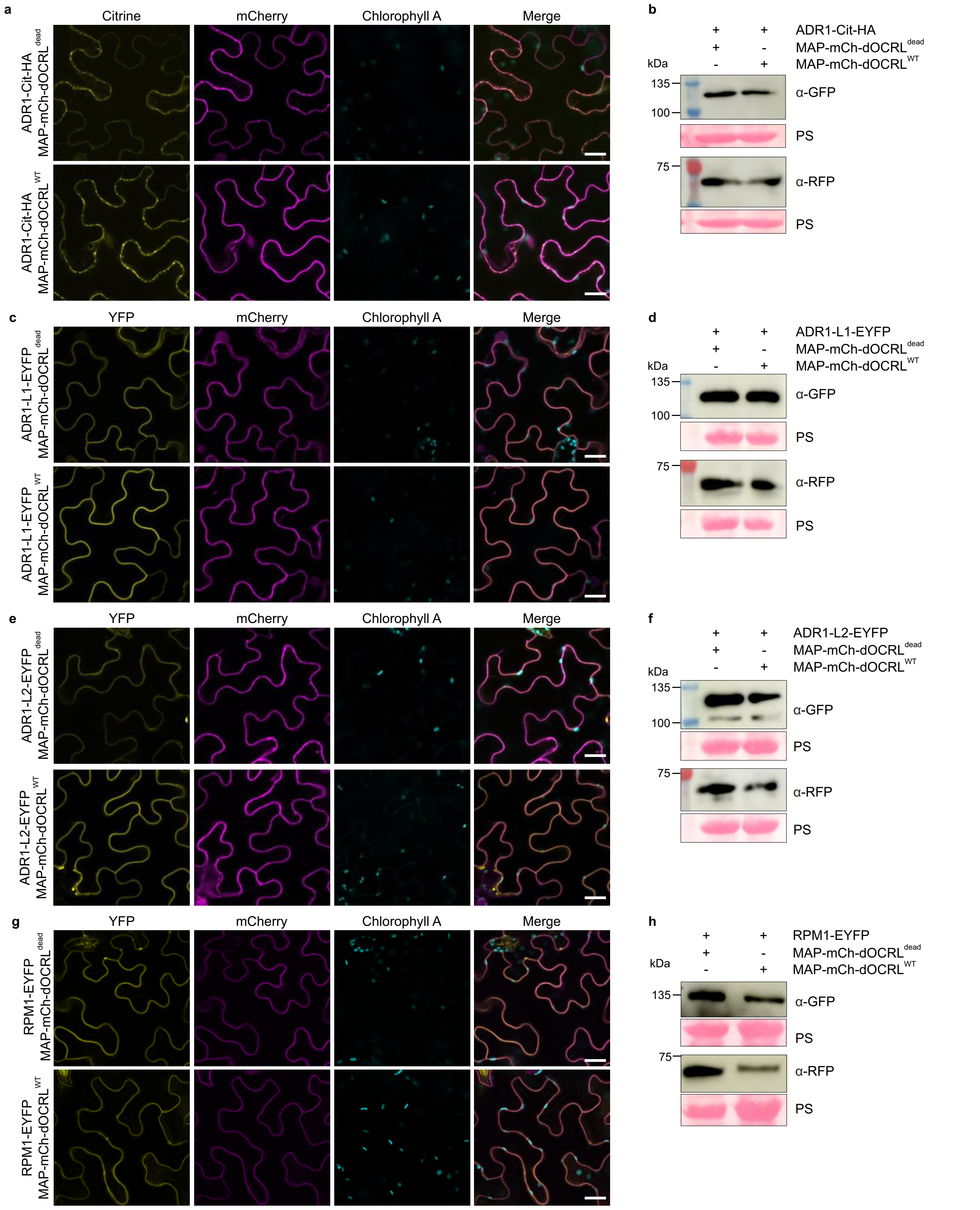

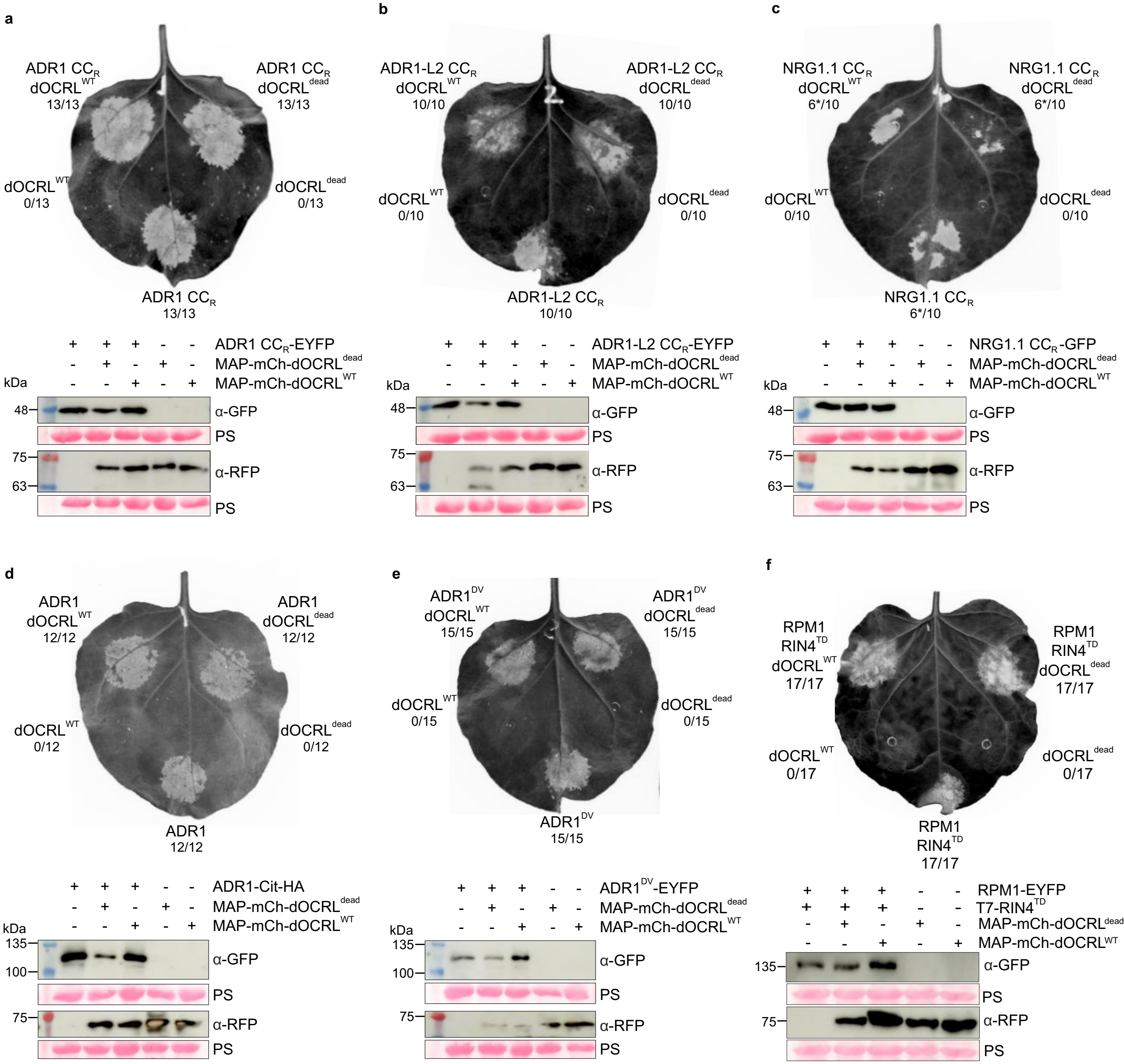

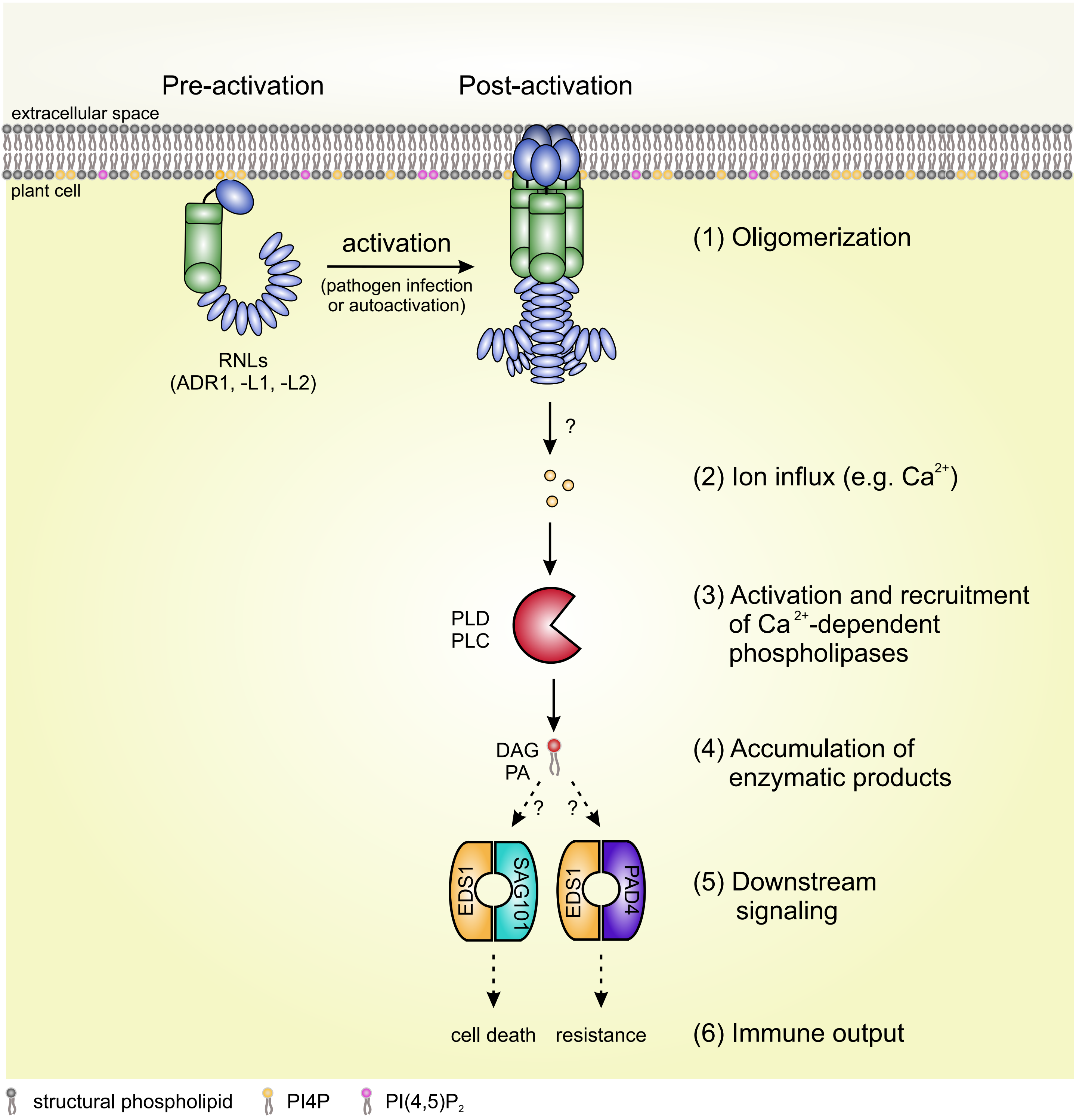
